# Denaturation resistant P2 tetramer is required to import fatty acids into intraerythrocytic Plasmodium falciparum

**DOI:** 10.1101/2022.06.11.495734

**Authors:** Sudipta Das, Kaustav Majumder, Anwesha Bhattacharjya, Bhaskar Roy

## Abstract

The initiation of asymmetric karyokinesis of intraerythrocytic Plasmodium falciparum (Pf) begins without dismantling the nuclear envelope showing the hallmark feature of closed mitosis (1-10). In Pf, karyokinesis precedes cytokinesis and cell body formation (6, 8-10). Regulation at the beginning of nuclear division either through checkpoints or by importing serum components was largely unknown. At the trophozoite stage, PfP2 tetramer trafficked to the infected erythrocyte (IE) surface and the inaccessibility of IE surface PfP2 to its bonafide ligand led to the arrest of nuclear division (11-13). Here we show that PfP2 tetramer localization on the IE surface and the beginning of nuclear division are concomitant in nature. Synthetically induced denaturation resistant PfP2 tetramer interacts with human serum fatty acids and phospholipids for its import into IEs at the beginning of karyokinesis. In the natively folded denaturation resistant PfP2 tetramer cage, the Cys-Cys redox switch regulates the binding and subsequent release of fatty acids on the IE surface. This mechanistic insight of fatty acids import inside IEs using synthetically induced denaturation resistant PfP2 tetramer provides an unique drug screening platform for novel small molecule screening against malaria.

## Introduction

Plasmodium falciparum (Pf), the causative agent of malaria, remains a huge public health burden globally (14). During intraerythrocytic development in humans, Pf shows closed mitosis during schizogonic cell division where the parasite nucleus divides asymmetrically often showing an odd number of nuclei in the shared nucleoplasm without dismantling the nuclear envelope. Chromosomes of a replicating parasite get segregated towards MicroTubule Organising Centre (MTOCs) at the opposite pole without dismantling the nuclear envelope giving rise to two daughter nuclei without cytokinesis. The asymmetric number of daughter nuclei increases within the shared nucleoplasm keeping nuclear envelop intact showing hallmark features of closed mitosis (1-8). In addition to protozoan parasites, nuclear asymmetry has been observed in many primitive unicellular eukaryotes such as fungi (15), insects (16). In these organisms, how the asymmetry is being achieved and maintained is not clear. Are there any roles of checkpoints before the initiation of asymmetric karyokinesis that need to be further investigated. In intraerythrocytic Pf, the molecular mechanism of karyokinesis and reasons for autonomous nuclear division in a pool of asymmetric nuclei are poorly understood. However, autonomy in the nuclear division has been speculated to be due to local skewing of nutrients or might be due to limiting factors that slows down the multiplication of nuclei as it progresses towards mature schizogony (5-10),. In intraerythrocytic Pf, Checkpoint(s) at the beginning of karyokinesis or before the commitment of repeated DNA synthesis and mitosis (S/M phase) might play a significant role in successfully completing the energy and resource requiring S/M phase (17-20). But before karyokinesis or during the ongoing S/M phase when and how these checkpoints in intraerythrocytic Pf are operational are still largely unknown and need to be pondered to understand the fundamental biology as to how karyokinesis begins and the molecular cascade behind it.

During intraerythrocytic development of Plasmodium, several PEXEL and non-PEXEL effector proteins trafficked to the infected RBC (iRBC) surface which is predominantly virulence factors and channel proteins, playing important roles in disease biology and solute uptake, respectively (21-24). PfEMP1 and PSAC are the two well consensus virulence factors and channel proteins on the iRBC surface respectively (21-25). In addition, other PEXEL and non-PEXEL proteins have been reported to translocate to the iRBC surface which renders the iRBC membrane more promiscuous and nonselective to solutes and other macromolecules towards the trophozoite stage and schizogony (26-27). Strangely, 60S stalk ribosomal protein P2, being a non-PEXEL protein has been reported to be present on the iRBC surface during the mid-late trophozoite stage (11-13). Post-merozoite invasion (PMI) at around 30 hrs, P. falciparum P2 (PfP2) was found to be present on the iRBC surface only for 3-4 hrs, that to as a DTT/β-ME, SDS, and boiling resistant tetramer (11-13). When iRBC surface PfP2 was made unaccessible to its bonafide ligand(s) through monoclonal antibody binding, karyokinesis of Plasmodium was arrested at the di-nucleated trophozoite stage at the very beginning of nuclear division^11^. In those arrested parasites, tubovesicular network (TVNs) and lipid imports were significantly diminished may be due to the non-functional PfP2 tetramer which was made unaccessible to the ligand through antibody blockage (11). In other apicomplexan parasites, e.g., *Toxoplasma gondii* (Tg), P2 was localized on the tachyzoite surface and implicated in the invasion of the host cells (28). In yeast, P2 was localized on cell surface either as a monomer or as a heteromeric protein complex (29). In Neisseria gonorrhea (Ng), the orthologue of P2 is L12, which was reported to be localized on the bacterial cell surface and implicated in host cell invasion (30).

In solution, recombinant PfP2 (rec.PfP2) forms a molten globule tetramer having hydrophobic pocket on the surface (31,32). NMR clearly showed that rec. PfP2 protein has a high propensity to oligomerize and tend to form oligomers at physiological pH and temperature (33,34). Circular dichroism (CD) and solution NMR has also revealed that the monomeric rec. PfP2 is predominantly α helical but molten globule in nature and the “C” terminal region is intrinsically disordered (33,34). 2D ^1^H–^15^N HSQC spectra of tetrameric rec. PfP2 exhibited hydrophobic surface mostly contributed by the N terminal α helices as shown using 8-anilinonaphthalene-1-sulfonic acid (ANS) (31), indicating that rec. PfPP2 tetramer could provide sites for intermolecular association in an aqueous environment and may have the possibility to bind non-polar/hydrophobic molecules as a bonafide ligand(s) on iRBC surface. In a recent discovery, it has been demonstrated that rec.PfP2 tetramers stabilise themselves on the iRBC surface by interacting with Band 3 protein where N terminal 70 amino acids of PfP2 interact to form the oligomers and associate with Band 3 protein (35). Between the *P. falciparum* P2 and the human P2, there is 69% amino acid sequence homology but they differ in their oligomerization pattern and behaviour as the human P2 at physiological pH forms a stable dimer (36) but under same condition, *P. falciparum* P2 forms a molten globule tetramer indicating critical functional implications of tetrameric P2 on the iRBCs surface during mid-late trophozoite stage.

Unlike other apicomplexan parasites, localization of PfP2 tetramer to the host cell surface concomitant with the initiation of karyokinesis drove our attention towards its possible role in the regulation of nuclear division possibly by acting as a gatekeeper / importing complex on the iRBC surface. We hypothesized that PfP2 tetramer either independently or in association with other parasite proteins form a homo/hetero complex which might be involved in sensing household before karyokinesis begins or might act as a non-polar or hydrophobic molecule importing complex critically required for the nuclear division to proceed. Hence blocking tetrameric PfP2 on the iRBC surface led to the arrest of nuclear division (11,12). This astonishing previous discovery posed a genuine question as to what is/are the bonafide ligand(s) of PfP2 tetramer on the iRBC surface blocking of which led to the arrest of karyokinesis. To identify that ligand(s) from human serum and to understand the nature of denaturation resistant PfP2 tetramer on the iRBC surface, we have performed a plethora of biochemical, biophysical experimentations to reveal the function of PfP2 tetramer on the iRBC surface and its mechanism of binding/release of ligand(s) towards the discovery of a novel anti-malarial drug target during intraerythrocytic development in human erythrocytes.

## Results

### Localization of PfP2 tetramer on the iRBC surface and the beginning of intraerythrocytic karyokinesis are concomitant

At the early-mid trophozoite stage, 60S stalk ribosomal P2 protein is trafficked to the infected RBCs (iRBCs) surface. Out of the monomeric and tetrameric pool of P2 protein in the parasite cytosol, only tetramer species seem to get exported to the iRBCs surface for 3-4 hrs (11,12). Blocking the accessibility of PfP2 using specific monoclonal antibodies or down regulation of PfP2 resulted in nuclear division arrest of the parasites at the very beginning of karyokinesis (11,12) (Supplementary data fig.2b & c). After 3-4 hrs of PfP2 localization on the iRBCs surface (Supplementary data fig.2d & e), it disappeared from the surface as IFA did not catch any fluorescence signal (11). The iRBCs surface localized PfP2 tetramer appears to be DTT and SDS resistant as the iRBCs ghost showed the presence of tetramer PfP2 which did not resolve to a monomer under reducing SDS-PAGE (11-13). To understand the nature of PfP2 tetramer on the iRBCs surface and its critical role at the beginning of parasite karyokinesis, in vitro we reconstituted denaturation resistant recombinant PfP2 (rec.PfP2) tetramer. PfP2 was recombinantly expressed and purified in E. coli as a 6X His-tag protein (Supplementary data fig. 1). Purified rec. PfP2 has a propensity to form oligomers when incubated at 37°C at physiological pH. Under the native condition, rec.PfP2 distinctly forms monomer, dimer, and tetramer (Supplementary data fig.1a, b & e). Immunoblot using rabbit polyclonal sera generated against rec. PfP2 did bind all the species of rec. PfP2 depicts the inherent nature of oligomerization under the non-reducing conditions in SDS-PAGE (Supplementary data fig.1c). When trophozoite stage parasite lysate was probed using rabbit polyclonal antibody (10k dilution), parasite P2 protein was observed as a monomer and tetramer showing the specificity of the rabbit sera to PfP2 (Supplementary data fig.1d).

To understand the concomitant nature of the localization of PfP2 to the iRBCs surface and the initiation of parasite karyokinesis, PfP2-HA transgenic parasites (Supplementary data fig. 2) were synchronized and at around 18-22h post-merozoite invasion (PMI), taxol was added in parasite culture medium (Fig. 1a). After 6h of treatment, karyokinesis arrested iRBCs were stained for the localization of PfP2 and the nucleus (Fig. 1a & b). Immunofluorescence assay (IFA) and imaging through confocal optical slices of taxol arrested iRBCs distinctly showed the punctate nature of PfP2 on the iRBCs surface and in the iRBCs cytosol. Around 90-95% taxol arrested transgenic parasites showed PfP2 on the iRBC surface (Fig. 1c & d). The arrested parasite nucleus appears to be doublet in nature when stained with DAPI (Fig. 1b). At 28-30h PMI at the trophozoite stage, around 85-90% of transgenic parasites showed PfP2-HA on the iRBC surface (Supplementary data fig. 2d & e) aligning with the fact that the localization of PfP2 tetramer on the iRBC surface and the beginning of karyokinesis is a tightly coupled event possibly involved in determining the progression of karyokinesis and cell body formation by importing serum components through PfP2 tetramer complex on the iRBC surface. The cytosol of wild type and transgenic PfP2-HA parasites clearly showed two species of the protein, monomeric PfP2 and denaturation resistant PfP2 tetramer (Fig. 1e). In the presence of 3 mM glucosamine, cytosolic PfP2 was diminished significantly resulting in the arrest of parasite karyokinesis (Fig. 1f, Supplementary data fig. 2b & c). Immunoprecipitation (IP) of iRBC ghost from wild type and transgenic parasites using PfP2 rabbit polyclonal and anti-HA antibody revealed the presence of denaturation resistant PfP2 tetramer on iRBC surface (Fig. 1g & h) suggesting a molecular event occurring in the trophozoite stage parasite cytosol resulting in the formation of denaturation resistant PfP2 tetramer without the involvement of S-S bond of cysteine residues and subsequent trafficking of the tetramer to the iRBC surface.

**Fig. 1.**
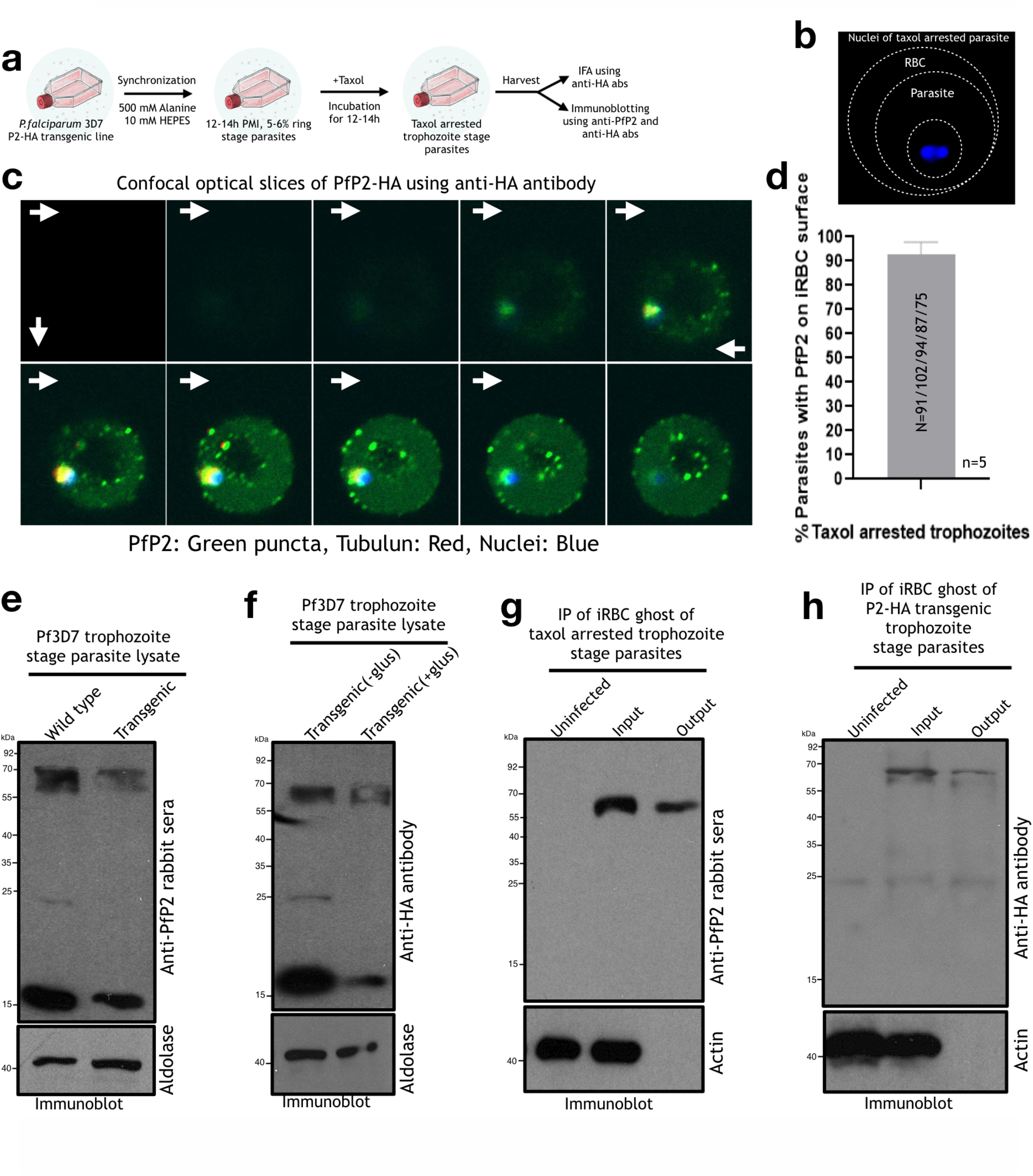
Initiation of parasite karyokinesis and localization of PfP2 tetramer on the IE surface are concomitant. **a**. Schematic depicting the process of synchronization and taxol treatment of P. falciparum P2-HA transgenic parasites and subsequent usage of taxol arrested parasites. **b**. Representative image of a taxol arrested intraerythrocytic parasite nuclei at trophozoite stage. **c**. Immunofluorescence assay (IFA) of taxol arrested transgenic P2-HA P. falciparum 3D7 parasites followed by confocal optical sectioning through Z axis. Parasite P2 (green puncta) was stained with anti-HA antibody, parasite β tubulin (red) was satined with anti-β tubulin antibody. Parasite nuclei was stained with DAPI. White arrow depicts the initiation of optical slice and the subsequent revelation of parasite P2 protein (green puncta) at every Z stack. **d**. Quantification of taxol arrested P2-HA transgenic parasites at trophozoite stage showing P2 protein on infected RBC (iRBCs) surface. Five biological replicates of taxol treated parasites showed around 90-95% arrested nuclei at trophozoite stage where P2 protein is present on the iRBC surface. (n=5). N represents the total number of parasites counted for arrested nuclei and P2 protein on the iRBC surface. **e**. Immunoblot of wild type and transgenic P. falciparum 3D7 trophozoite stage parasites. Wild type parasite lysate (2 μg) and transgenic parasite lysate (1.4 μg) was separated in reducing SDS-PAGE and probed with anti-PfP2 rabbit serum. Parasite aldolase was used as a loading control. **f**. Immunoblot of transgenic trophozoite stage parasites in the absence/presence of glucosamine (3 mM). Around 1.5 μg of parasite lysate was separated in reducing SDS-PAGE and probed with anti-HA antibody. Parasite aldolase was used as a loading control. **g**. Immunoprecipitation (IP) of iRBC ghost of taxol arrested wild type parasites using anti-PfP2 rabbit polyclonal antibody. Total of 4 μg of iRBC ghost protein (input) was incubated with 8 μg rabbit polyclonal anti-PfP2 IgG. IP material was separated in reducing SDS-PAGE and immunoblotted using anti-PfP2 rabbit polyclonal antibody. As a control, uninfected RBC ghost total protein (4 μg) was used to check any antibody cross reactivity. Ghost actin was used as a loading control. **h**. Immunoprecipitation (IP) of iRBC ghost of taxol arrested P2-HA transgenic parasites using anti-HA antibody. Total 3 μg of iRBC ghost protein (input) was incubated with 6 μg anti-HA immunoglobulin molecules. IP material was separated in reducing SDS-PAGE and immunoblotted using anti-HA antibody. As a control, uninfected RBC ghost total protein (4 μg) was used to check any HA antibody cross reactivity. Ghost actin was used as a loading control. All immunoblot images were representative and the experiment and corresponding findings were repeated > 5 times.

In yeast, the P2 knockout strain did not show any growth defects, and P. falciparum P0 protein when complemented in yeast, did not require PfP2 or yeast P2 for ribosomal activity (37). These studies clearly suggested that PfP2 may not be essential for ribosomal activity but when the gene was attempted to be knocked out, it was found to be refractory to deletion during intraerythrocytic development^11,38,39^ confirming that the localization of PfP2 on the iRBCs surface is critically important during trophozoite stage, blocking of which either by antibody or through genetic knockdown inhibited/reduced PfP2 tetramer interaction with its bonafide ligand(s), hence did not allow parasites to proceed through schizogony. This observation posed a genuine question as to what is/are the serum component(s) binding to PfP2 tetramer on the iRBC surface.

### Synthesis of denaturation resistant iRBC ghost PfP2 tetramer and finding a needle in a haystack

On the iRBCs surface, PfP2 tetramer exists as a denaturation resistant complex which did not resolve to a monomer even after boiling in DTT treated SDS-PAGE (11-13). It was difficult to acquire sufficiently pure iRBC ghost tetrameric PfP2 material required for subsequent biochemical and biophysical experimentations to fish out the reasons for denaturation resistance. Using a unique approach, *in vitro* we successfully synthesized denaturation resistant rec.PfP2 tetramer where rec. PfP2 was incubated with Albumax (a parasite culture component rich in lipids and bovine serum proteins). But rec. PfP2 was not incubated with complete albumax, instead, albumax was dissolved in 1xPBS and then passed through a 5 kDa membrane cutoff filter which did not allow serum protein components in the passed through solution. Hence, after cutoff filtration, we used **L**ower **A**lbumax **F**raction **S**olution (LAFS) which predominately possessed small molecules present in the albumax and was mostly devoid of serum proteins (Fig. 2a). When rec. PfP2 was incubated with LAFS at 37°C for 3h and resolved in non-reducing SDS-PAGE, all the species (monomer and dimer) of rec. PfP2 converted into a stable tetramer (Fig. 2a & b). Whereas 6x His tag control proteins from mammalian and plant origins did not show any such transformation when incubated with LAFS (Fig. 2c), suggesting a specific ligand binding to rec.PfP2 and subsequent transformation to a tetramer. After this surprising and thought provoking observation, we wondered how fast this transformation is happening from monomer and dimer to a tetramer. To resolve this, rec. PfP2 was incubated with LAFS at 37°C for varying times and resolved in non-reducing SDS-PAGE. Within 30 mins of incubation, all the species of rec. PfP2 was transformed into a stable tetramer (Additional Supplementary Fig.1a). We were puzzled to observe this fast transformation of rec. PfP2 into a stable tetramer and tried to interrogate what could be the driving force behind it. At this stage we hypothesized that small molecule(s) could be of any nature present in LAFS has the potential to drive rec. PfP2 oligomerization into a tetramer. To understand the authenticity of this transformation, we simply titrated out small molecule(s) from LAFS by incubating rec. PfP2 with LAFS followed by affinity purification with Ni-NTA beads which binds 6X His tag in the rec. PfP2 tetramer. LAFS induced rec.PfP2 tetramer bound to Ni-NTA beads due to 6X His tags hence after centrifugation most of the rec. PfP2 molecules came out in the beads and supernatant was devoid of rec. PfP2 and the small molecules required for the induction of oligomerization. In that second supernatant of LAFS, when rec. PfP2 having monomers, dimers was incubated again for 3 hrs at 37°C, to our surprise, tetramerization was diminished significantly and we observed monomer, dimer and low level of tetramer (Fig. 2d). It clearly suggested that in LAFS, the quantity of ligand(s) whatever it may be (possibly small molecules) was reduced due to its consumption when the first time rec. PfP2 was incubated hence during the second incubation of rec. PfP2 with one time used LAFS, enough small molecule (s) were not available for the second transformation to rec. PfP2 tetramer. How does LAFS induced rec. PfP2 tetramer behaves as compared to inherent rec. PfP2 tetramer and whether it shows DTT/beta-ME resistance like PfP2 tetramer in iRBCs ghost (11,12) was a thrilling question to address. When rec. PfP2 was incubated with LAFS for varying time intervals and treated with 100 mM beta-ME containing loading dye followed by SDS-PAGE without boiling, at 0 time point, monomer and tetramer were observed and at subsequent time points, monomer disappeared and only tetramer was visible (Additional Supplementary fig.1b). This clearly suggested a fast dynamics for LAFS induced tetramer formation. Rec. PfP2 tetramer on the other hand, when checked for its sensitivity to reducing agent, it did get reduced to a dimer and monomer as opposed to LAFS treated tetramer (Supplementary Data fig. 1b) which indicated that at the fundamental level molecular interactions amongst amino acids are quite different between these two types of tetramers. Further, we went ahead to check whether LAFS treated rec. PfP2 tetramer can be resolved to a monomer after boiling under the reduced condition in a SDS-PAGE. As evident repeatedly, the dimer of rec. PfP2 and LAFS treated PfP2, both are resistant to reducing agent,, whereas tetramer of rec. PfP2 came down to dimer and monomer after DTT/beta-ME. treatment. More interestingly, LAFS treated PfP2 tetramer, even after beta-ME /DTT treatment and boiling did not get resolved to a monomer but surprisingly dimer did (Additional Supplementary fig.1c). This clearly suggested that after LAFS treatment, PfP2 tetramer acquired some unique conformation through novel covalent interactions amongst amino acids and as a reflection of these interactions, tetramer became denaturation resistant. It also suggested that rec. PfP2 dimer in LAFS treated sample was formed just as inherent rec. PfP2 dimer which was beta-ME/DTT resistant but boiling sensitive. indicating that LAFS treated PfP2 tetramer did not form by the assembly of two dimers, instead, this tetramer plausibly form very uniquely by the assembly of PfP2 monomer through novel interactions amongst inter-peptide amino acids possibly covalently linked. To uncover this possibility of inter-peptide covalent linkage formation, we finally went ahead and checked the stability of LAFS treated PfP2 tetramer in 4M urea SDS-PAGE with boiling under reduced conditions (Fig. 2e). LAFS treated PfP2 tetramer at three timepoints, did not get resolve to a monomer however a faint band of monomer was observed due to the reduction of dimers into monomer after boiling. This is precisely the reason why in the same urea gel, under non-boiling conditions, in the lane of rec. PfP2 and at three time points of LAFS treatment, dimers were visible (Fig. 2e). When LAFS from human serum was incubated with rec.PfP2 at 37 for 3h and for varying time, rec.PfP2 transformed into a tetramer that was denaturants resistant including 4M urea (Fig. 2 f, g & h). All these observations strongly suggested that LAFS treatment led to the formation of an unique tetramer assembly possibly covalently linked amongst amino acids of four PfP2 peptides.

**Fig. 2.**
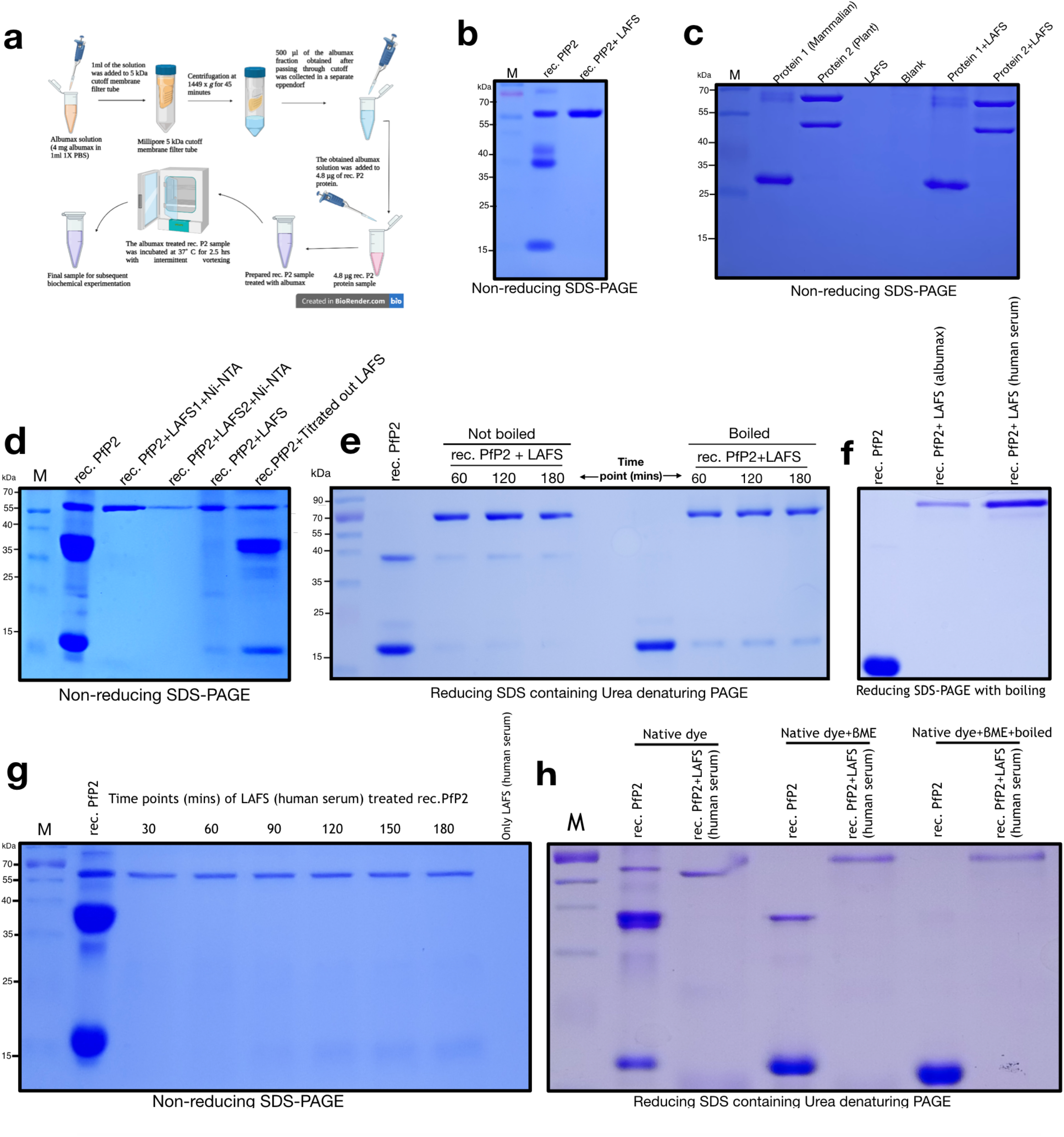
Components of human serum/albumax induce denaturation resistant PfP2 tetramer. **a**. Schematic showing the method of LAFS preparation and the treatment of rec.PfP2 with LAFS. **b**.0.5 μg of 6x His-tag rec. PfP2 was treated with LAFS for 3h at 37°C and separated in non-reducing SDS-PAGE. Rec. PfP2 in PBS for 3h at 37°C was used as a control. **c** 1 μg of 6x His-tag recombinant protein from mammalian and plant origin was treated with LAFS at 37°C for 3h and separated in non-reducing SDS-PAGE. Only LAFS was checked for any protein leftover from albumax solution. **d**. LAFS was titrated out with 1 μg of 6x His-tag rec.PfP2 at 37°C for 3h followed by affinity purification of LAFS treated rec.PfP2 tetramer using Ni-NTA beads. 1 μg of 6x His-tag rec.PfP2 was again treated with titrated out LAFS at 37°C for 3h and separated out in non-reducing SDS-PAGE. **e**. 0.5 μg of 6x His-tag rec. PfP2 was treated with LAFS for varying times (1h, 2h and 3h) and separated in SDS and 100 mM β-ME containing 4M Urea gel with/without boiling for 10 mins before sample loading. **f**. 0.5 μg of 6x His-tag rec. PfP2 was treated with LAFS from albumax and LAFS from healthy human serum at 37°C for 3h and separated out in reducing SDS-PAGE with boiling. **g**. 1 μg of 6x His-tag rec. PfP2 was treated with LAFS from human serum for varying times (30 mins-180 mins) at 37°C and separated in non-reducing SDS-PAGE without boiling. PBS treated 6x His-tag rec. PfP2 was used as a control. Only LAFS from human serum was loaded in the gel to check any leftover protein components in LAFS. **h**.0.5 μg of 6x His-tag rec. PfP2 was treated with LAFS from human serum at 37°C for 3h and separated in SDS and 100 mM β-ME containing 4M Urea gel with native loading dye (no SDS, no β-ME), with native dye+ β-ME but non-boiling and with reducing dye containing SDS, 100 mM β-ME and boiled. All the polyacrylamide gels were stained with coomassie R-250. All the gel images are the best representative and each type of experiment was repeated more than 4 times. M=Marker of protein ladder. LAFS: Lower Albumax Fraction Solution, collected from albumax and from a healthy human serum using 5 kDa membrane cutoff.

*Neisseria gonorrhoeae* (Ng) P2 ortholog exists at the bacterial cell surface and is implicated in host cell invasion (30). In Ng, Lysine-O-Cysteine (NOS) covalent bond mediated redox switch is one of the key regulators in many enzymatic activity (40,41). The widespread occurrence of NOS bond in many enzymes across eukaryotic species depicts its essentiality in the regulation of enzyme activity in general (40,41). Free radicals in LAFS might mediate NOS bond formation amongst four P2 molecules (41-43) hence we hypothesized that denaturation resistant PfP2 tetramer might be assembled by NOS covalent linkages where Lysine and cysteine residues in four P2 molecules are directly involved. In an experimental strategy, all cysteine residues in rec. PfP2 molecules were pre-reduced and then alkylated with Iodoacetamide (IAA) to prevent NOS bond formation (Supplementary Data fig. 3a). When LAFS was incubated with this modified rec.PfP2 molecules, we still observed denaturation resistant rec.PfP2 tetramer but interestingly, monomer to tetramer conversion was reduced hence intense monomeric bands were visible as opposed to without pre-reduction (Supplementary Data fig. 3b). These alkylated LAFS induced rec.PfP2 tetramers were β-ME, boiling and urea resistant confirming again that LAFS induced rec.PfP2 tetramer is formed due to unknown covalent linkages other than NOS (Supplementary Data fig. 3c & d).

On the iRBCs surface, whether PfP2 exists as a tetramer or does it form a higher homogeneous complex induced by acyl chains of RBC membrane resident lipids was a critical question to address. Since iRBCs ghost PfP2 showed SDS resistance, we intentionally chose SDS as a surfactant that possesses saturated acyl chain-like membrane surfactant lipid molecules e.g., phosphatidylcholine, phosphatidylserine etc (Supplementary Data fig. 4a). We wanted to check whether the acyl chain by itself has any effect on rec, PfP2 oligomerization/degradation when SDS concentration was used below CMC, at CMC and above CMC (Critical Micelle Concentration). The main provocation to understand the effect of acyl chain on the oligomerization and the stability of PfP2 tetramer/higher complex on the iRBCs surface came when IFA to visualize PfP2 tetramer on the iRBCs surface did not show any fluorescence after 36h PMI concomitant to the completion of its function (11,12). This observation, drove our attention towards a possible mechanism of self disassembly of PfP2 tetramer from the iRBCs surface after its critical function is over at the beginning of karyokinesis. Rec. PfP2 in blue Native PAGE showed an inherent propensity to form oligomers. Astonishingly, oligomerization was aggravated in the presence of SDS and tetramer and higher oligomers showed DTT/beta-ME sensitivity whereas dimer was unaffected under reducing conditions and did not resolve to a monomer (Supplementary Data fig. 4b). This seems to suggest that the mechanism of dimer formation differs from the mechanism of tetramer formation. It is indeed an unique example, where the same protein behaves differently at different oligomerization state up to an extent where tetramer seems to get stabilized by disulfide bridges wheres dimer is built through interactions other than disulfide linkages. It is still not clear whether rec.PfP2 tetramer is formed directly due to the association of monomers or two dimers assembled to form a tetramer. To understand the effect of the acyl chain of SDS on the oligomerization/dissolution of rec. PfP2 tetramer, we designed blue native PAGE, where rec.PfP2 was incubated with a gradient concentration of SDS starting from below CMC to above CMC. Rec. PfP2 tetramer was distinctly visible at the SDS concentration just before CMC and at CMC, but surprisingly, at above CMC the rec. PfP2 tetramer self-dissociate as it is evident because the pixel density of coomassie stained tetramer diminished significantly compared to the concentration at CMC (Supplementary Data fig. 4d). Whereas LAFS treated rec.PfP2 tetramer remain as a stable species even after SDS treatment at any concentration (Supplementary Data fig. 4c). When other surfactants which did not have acyl chains, such as CHAPS, Na-deoxycholate and Triton-X100 were used at different gradient concentrations and treated with rec.PfP2, neither they induced the propensity of tetramer formation nor at any concentration the inherent rec. PfP2 tetramer self-dissociated (Supplementary Data fig. 4e, f & g). This seems to suggest a critical role of the acyl chain of RBC resident lipid surfactant molecules in the assembly and/or dissolution of PfP2 tetramer complex on the iRBCs surface. It is still elusive as to how the above CMC concentration of SDS triggers dissociation of rec. PfP2 tetramer and how it is being achieved on the iRBC surface after the function of the tetramer complex is over. To visualize the structure of rec. PfP2 tetramer using Atomic Force Microscopy (AFM), inherent and LAFS treated tetramer species were isolated by size exclusion chromatography (Supplementary Data fig.5a & b). rec.PfP2 population consisting of monomer, dimer and tetramer and isolated tetramer showing distinct structural conformation where one tetramer molecule is formed by S-S linkages of cysteine residues between two dimers (Supplementary Data fig. 4b, 5d). The width of rec.PfP2 tetramer was 130 nM and the passage between two dimers is 4-5 nM (Supplementary Data fig. 5d). At low magnification, many tetramer particles showed similar conformation indicating a mechanism driven tetramer formation. LAFS (from albumax) and SDS treated rec.PfP2 tetramer showed entirely different structural conformation where unlike control tetramer, LAFS and SDS treated tetramer looks more circular without any visible passage between two dimers suggesting conformational flexibility between ligand bound/unbound state (Supplementary Data fig. 5c & d). Mass of rec.PfP2 tetramer and LAFS treated rec.PfP2 tetramer was found to be ∼66 kDa, whereas only LAFS did not show protein mass peak in that range corroborating with the coomassie gel finding showing no protein band (Supplementary Data fig. 6a & b). Inspite of having closely similar molecular weight, LAFS treated rec.PfP2 tetramer was more compact and smaller in hydrodynamic radius as compared to rec.PfP2 tetramer (Supplementary Data fig. 6c & d). In Blue Native PAGE, LAFS treated rec.PfP2 tetramer migrated much faster (Supplementary Data fig. 6e). This suggested that after LAFS treatment, rec.PfP2 tetramer acquires a more ordered and compact conformation plausibly required to interact with and hold serum components for its importing activity on the iRBC surface.

### PfP2 tetramer on the iRBC surface serves to import fatty acids

The malarial parasite has an enormous requirements for fatty acids during its replicative cycles in mammalian host. Salvage through import and de novo biosynthesis of fatty acids both are operational in malaria parasites depending on type of mammalian cells are infected and supporting parasite’s replicative cycles. Intraerythrocytic (IE) development is solely depended on imports of serum fatty acids (44-46) whereas during liver and mosquito stages, FAS type II pathway in the parasite apicoplast is responsible for de novo biosynthesis (44-49). The FAS-II pathway is responsible for the elongation of fatty acids via the action of four distinct enzymes: FabG, FabZ, FabI, and FabB/F (50-52) When these enzymes were deleted from blood stage parasites, effect on growth rate was minimal strongly suggesting an active fatty acids import mechanism from serum through a protein complex on iRBC surface. At the beginning of intraerythrocytic schizogony, import of serum fatty acids is critically important for new membrane biogenesis hence blockage of import leads to shortage of one of the household which is necessary to carry forward karyokinesis (53-56). This checkpoint like event through a iRBC surface protein complex has been shown to be critical at the beginning of nuclear division concomitant with the localization of PfP2 tetramer on the iRBC surface.

To resolve what does denaturation resistant PfP2 tetramer on the iRBC surface bind to, rec.PfP2 was incubated with LAFS. Prepared LAFS (Fig. 3a) was subjected to ESI-MS (direct infusion) to understand the mass and nature of molecules present in LAFS (Fig. 3b). LAFS induced rec.PfP2 tetramer (Fig. 3c) was subjected for mass spectrometric analysis to detect LAFS components which were imprisoned in natively folded PfP2 tetramer cage. LC-MS profile of LAFS induced rec.PfP2 tetramer at different retention time distinctly identified many peaks corroborating adduct masses of fatty acids and surfactant phospholipids. These peaks were majorly absent in the control panel where inherent rec.PfP2 tetramer was subject for LC-MS without any LAFS treatment (Fig. 3d, e & f).

**Fig. 3.**
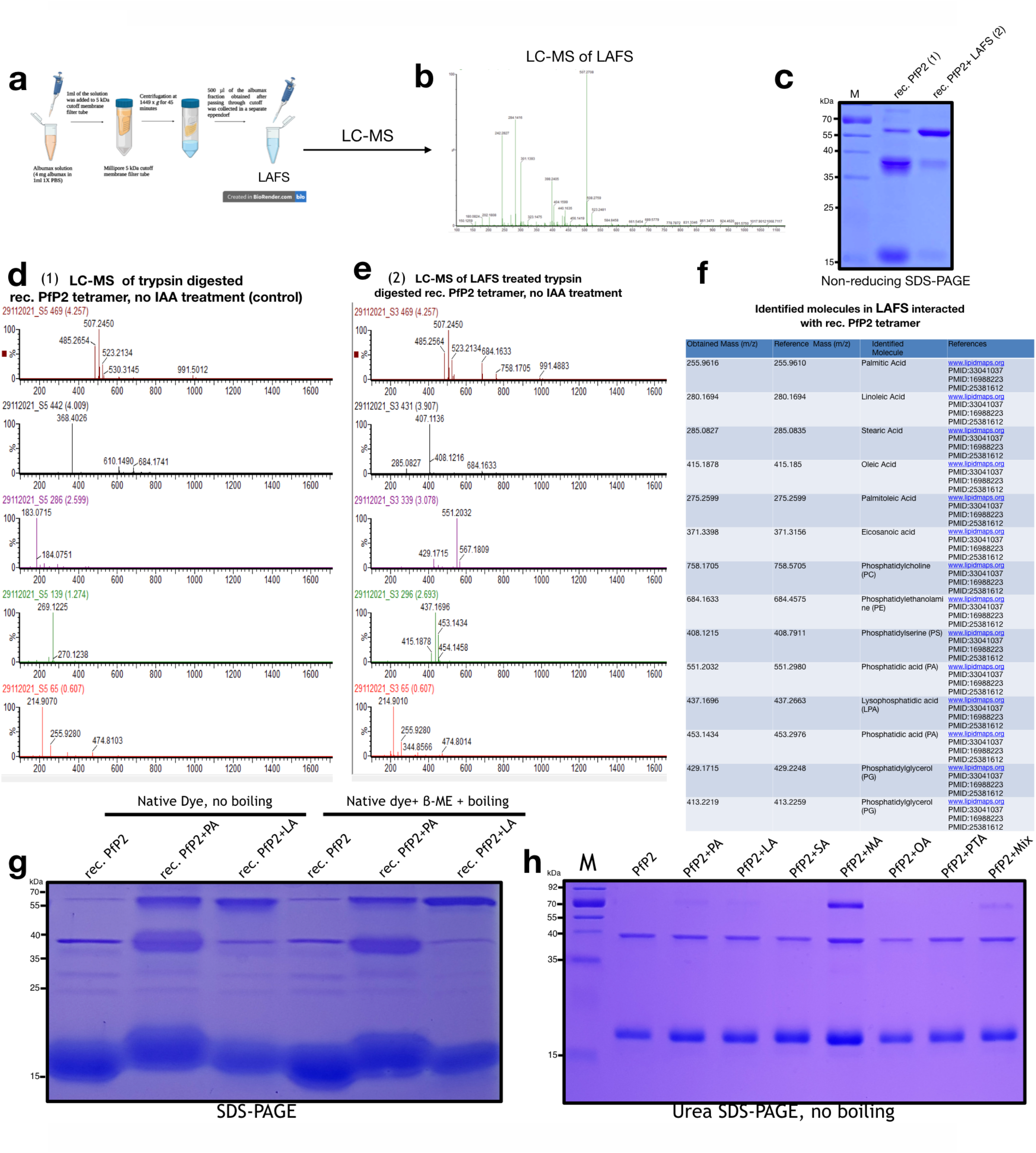
Denaturation resistant PfP2 tetramer binds fatty acids on infected erythrocyte surface. **a**. Schematic showing the method of LAFS preparation. **b**. LC-MS (direct infusion) of LAFS showing the mass of molecules present in LAFS at each peak. **c**. 0.7 μg of 6x His-tag rec. PfP2 was treated with LAFS for 3h at 37°C and separated in non-reducing SDS-PAGE. Rec. PfP2 in PBS for 3h at 37°C was used as a control. **d**. LC-MS profile at a different retention times of trypsin digested but not DTT and Iodoacetamide treated tetrameric rec, PfP2 (control) showing the different mass of P2 peptides which were generated from natively folded PfP2 tetramer. **e**. LC-MS profile at the different retention times of LAFS treated rec. PfP2 tetramer which was trypsin digested but not DTT and Iodoacetamide treated. LC-MS profile depicting different masses unrelated from panel c. LC-MS profile of control PfP2 tetramer and LAFS treated PfP2 tetramer were repeated > 10 times to check the reproducibility of the data. In the LC-MS profile, X axis represents m/z and Y axis represents % abundance. **f**. List of molecules in LAFS interacted with PfP2 tetramer identified by calculating adduct mass of each molecule found in panel e. To identify molecules using their adduct mass, www.lipidmaps.org was mainly used. **g**. 0.5 μg of 6x His-tag rec. PfP2 was treated with Palmitic Acid (PA) and Linoleic Acid (LA) at 10 mM concentration for 3h at 37°C and separated in SDS-PAGE. PfP2 in PBS for 3h at 37°C was used as a control. **h**. 0.5 μg of 6x His-tag rec. PfP2 was treated with Palmitic Acid (PA), Linoleic Acid (LA), Stearic Acid (SA), Myristic Acid (MA), Oleic Acid (OA), Palmiteloic Acid (PTA) and Mixture of all fatty acids (Mix) at 10 mM concentration for 3h at 37°C and separated in reducing Urea (4M) SDS-PAGE. All the polyacrylamide gels were stained with coomassie R-250. All the gel images are the best representative and each type of experiment was repeated more than 3 times.

When six different fatty acids were checked individually for its ability to interact and induce rec.PfP2 tetramer, all of them induced denaturation resistant PfP2 tetramerization with a varying degree. When Palmitic acid and Linoleic acid were incubated with rec. PfP2 tetramer and subjected to SDS-PAGE with/without boiling of samples, induced PfP2 tetramer did not resolve to a monomer corroborating with the reason for denaturation resistant PfP2 tetramer formation after LAFS treatment (Fig. 3g). When six individual fatty acids and a mixture induced PfP2 tetramer was separated in urea PAGE, with respect to untreated PfP2 control, varying degree of urea resistant tetramer was observed with an exception of myristic acid where tetramer population was significantly higher compared to other fatty acids (Fig.3h). These confirmed that PfP2 tetramer interacts with fatty acids and as a result acquires denaturation resistant conformation (Supplementary Data Fig. 5e).

Denaturation resistant PfP2 tetramer in iRBC ghost interacts with fatty acids. Mechanistically in this interaction, oxidation/reduction (redox switch) of cysteine residues in PfP2 tetramer appears to be involved in the binding and release of fatty acids. To explore this hypothesis, rec.PfP2 was pre-reduced with DTT and then alkylated using iodoacetamide (IAA) followed by LAFS treatment. When LAFS induced alkylated PfP2 tetramer was subjected for LC-MS to detect whether it can bind fatty acids, astonishingly none of the peak of adduct masses matched with any fatty acids, suggesting the involvement of cysteine residues in PfP2 tetramer for its interaction with serum fatty acids (Supplementary Data fig. 7c). Cys-Cys redox switch appears to regulate fatty acid binding and its subsequent release to maintain import activity of ghost PfP2 tetrameter complex on the iRBC surface.

## Discussion

At the beginning of karyokinesis, at the di-nuclear stage, PfP2 was found to be localized on the iRBC surface as a denaturation resistant tetramer (11,12). The timing of PfP2 tetramer localization on the iRBC surface and the initiation of nuclear division hinted at the concomitant nature of these two cell biological phenomena during intraerythrocytic karyokinesis. Taxol is a stabilizer of the microtubule and protects it from disassembly, arrested karyokinesis at the first nuclear division by inhibiting chromosome segregation and daughter nuclei formation concomitantly having PfP2 tetramer on the iRBC surface. This signified a critical role of iRBC surface PfP2 tetramer at this stage either as a checkpoint or as a nutrient (serum components) importing complex required to check and maintain household before the full commitment of cell division.

Import of small molecules and ions from serum occurs through PSAC, a well consensus anion channel on the iRBC surface (25). The selective removal of palmitic acid and oleic acid from the parasite culture media resulted in the arrest of parasite nuclear division at the same stage as anti-PfP2 antibody or PfP2 knockdown parasite showed (53-56). This clearly indicates that PfP2 tetramer on the iRBC surface serves in the import activity of serum fatty acids and other lipids during early schizogony. However, after binding and subsequent release of fatty acids into the iRBC cytosol, how it is being transported into the parasite is still not clear. To transport fatty acids in the parasite cytosol, either another protein complex is engaged or P2 oligomers on the parasite plasma membrane might be operational or it could be a simple diffusion across the parasite membrane. The tachizoite surface of Toxoplasma possesses P2 and it has been implicated for host cell invasion (28). It is not clear but it is likely that the Plasmodium plasma membrane also has P2 oligomers through which fatty acids are getting incorporated into the parasite cytoplasm (Fig. 4). P2 being a high expressing ribosomal protein, using either microscopy or membrane fractionation, it was difficult to distinguish whether the parasite membrane has P2 oligomers. Denaturation resistant PfP2 tetramer has been observed in the parasite cytosol after 24h PMI and then trafficked to the iRBC surface (11,12). Hence while crossing the parasite membrane and PVM, a subpopulation of PfP2 tetramer possibly retain themselves on to the parasite membrane for fatty acid import into the parasite. Either this retention of tetramer could be a mechanism driven process or it is a result of partial failure in successful trafficking of tetramer to the iRBC surface.

**Figure 4.**
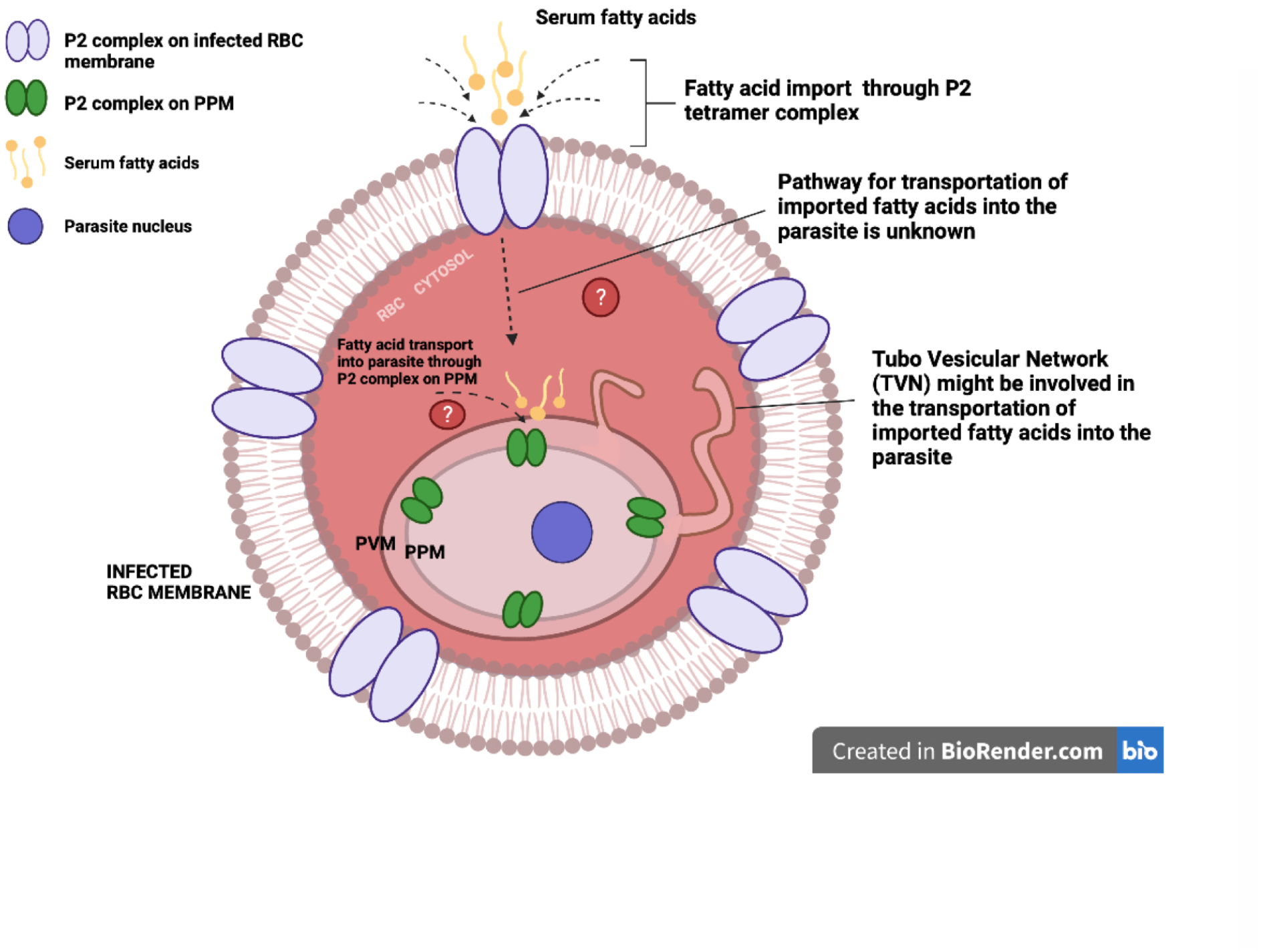
Model for fatty acids import through iRBC surface localised PfP2 tetramer complex. Model showing the distribution of denaturation resistant PfP2 tetramer (purple overlapped oval shaped) on an infected erythrocyte (IE) surface. Serum fatty acids import through the PfP2 tetramer complex IE surface into the IE cytoplasm. Imported fatty acids transport across IE cytoplasm either through diffusion or by an unknown transporter. As shown in toxoplasma, P2 exists on the surface of tachizoite (28). Parasite plasma membrane might also have P2 oligomers (green dumbbell shaped) and possibly be involved in the import activity of fatty acids from IE cytoplasm into the parasite for membrane biogenesis.

PfP2 interacts with fatty acids and appears to acquire denaturation resistant conformation. But when all cysteine resides were alkylated by IAA followed by LAFS treatment, PfP2 still formed denaturation resistant tetramer but it did not bind fatty acids, which clearly indicated that to acquire denaturation resistance, other possible covalent interactions four PfP2 monomers including fatty acids binding are required to achieve resistant conformation. PfP2 tetramer has 8 cysteine and 14 lysine residues. Four pairs of Cys-Cys redox switches in PfP2 tetramer appear to be involved in fatty acid binding and its subsequent release. Iodoacetamide treated PfP2 did form a denaturation resistant tetramer after LAFS treatment. But this tetramer did not bind serum fatty acids, strongly suggesting that irreversible changes in cysteine residues permanently block the fatty acid binding ability of the PfP2 tetramer. It is clear that cysteine residues must be free for them to be reversibly switched from a reduced state to oxidized state for fatty acid binding and its subsequent release. Iodoacetamide experiment clearly showed that under oxidized state of cysteine, fatty acids are binding and its subsequent release needs a reduction of cysteine moieties. It is still not clear how oxidation and reduction of cysteine residues are being achieved on the iRBC surface and whether serum components play any role in this type of switching activity. In addition to disulfide bond, cysteine residues are susceptible to oxidation by Reactive Oxygen Species (ROS), thus critically important for scavenging ROS and also required for cellular signalling and communications in different cellular contexts. Due to elevated ROS level at the trophozoite stage during hemoglobin degradation and hemozoine formation (57-59), functionally active PfP2 tetramer might form through Lysine-O-Cysteine (NOS) bridge as shown in the transaldolase enzyme of *Neisseria gonorrhoeae* (40,41). Oxidized cysteine being a highly reactive moiety, might also form a covalent linkage with tyrosine residue in the active site of the enzyme (60,61). PfP2 tetramer has four pairs of tyrosine and cysteine residues, hence ROS mediated Tyr-Cys covalent switch might also be involved in the binding and release of fatty acids on the iRBC surface. Overall understanding of how fatty acids are binding and released in the natively folded PfP2 tetramer cage would bring a mechanistic insight necessary for the development of better anti-malaria small molecules.

Phylogenetic analysis across many species (Supplementary data fig. 8) depicts that Plasmodium and human P2 have diverged, which seems to suggest that human P2 might only be involved in ribosomal activity whereas Plasmodium P2 depending on structural conformation might have additional non-ribosomal functions. In the apicomplexan clade, P2 is closely placed where parasites show rapid replication like schizogonic cell division. Toxoplasma P2 is distantly placed where cell division is symmetrical unlike Plasmodium. It appears, at least during schizogonic or rapid cell division process, P2 might have critical non-ribosomal functions which might be relevant in different cell biological contexts in many species hence warrant further in-depth investigations.

## Methods

### *P. falciparum* Parasite Culture

*P. falciparum* 3D7 parasites were cultured using type O^+^ human RBCs in RPMI1640 supplemented with 15 mM HEPES, 2 g/L sodium bicarbonate, 10 mg/L hypoxanthine, 50 mg/L Gentamicin sulfate, and 0.5% Albumax (cRPMI). Asexual stages of P. falciparum were maintained at 5% haematocrit in cRPMI at 37°C in a humidified incubator containing 90% N2, 5% O2 and 5% CO2. Parasite cultures were periodically tested by PCR for Mycoplasma contamination to ensure that they are free from Mycoplasma. Parasites were synchronised using 500 mM Alanine and 10 mM HEPES. Two rounds of synchronised 6-7% ring stage parasites were used for transfections.

### Treatment of *P. falciparum* infected RBCs with Taxol in culture

Paclitaxel (Sigma: T7191) was dissolved in DMSO to make a 10x concentration. Required volume from the stock was added into the culture medium to achieve a 500nM final concentration. P. falciparum parasites were synchronised 2 rounds using 500mM Alanine and 10mM HEPES. After 2 generations, highly synchronised ring stage parasites at around 16-18h PMI were subjected for taxol treatment. After 12-14h of treatment, at the trophozoite stage, at around 28-32h PMI, taxol arrested iRBCs were collected and washed twice with 1x PBS and used for biochemical experiments and confocal imaging.

### Immunofluorescence assay (IFA) and confocal imaging

IFA of taxol arrested iRBCs was performed in solution. Infected RBCs were centrifuged at 500*g* for 7 min, washed twice with 1xPBS, and re-suspended in PBS. Infected RBCs were fixed using 4% formaldehyde/0.0075% glutaraldehyde in PBS for 20 min at 4°C. All subsequent steps were carried out at room temperature (24–26°C). Infected RBCs were permeabiilzed using 0.1% Triton-X 100 in PBS for 30 min and washed three times with 1xPBS. 3% BSA in 1xPBS was used for blocking. Anti-HA antibody to PfP2-HA was used at 1:500 dilution in 1xPBS containing 0.01% Triton X-100 and incubated for 3–4h at room temperature. Infected RBCs were pelleted at 500x *g*, washed 3 times with 1xPBS containing 0.01% Triton X-100 and treated with appropriate Alexa 488 conjugated secondary antibodies (Molecular Probes) at 1:500 dilution for 2h at room temperature. After washing 3–4 times with 1xPBS containing 0.01% Triton X-100, iRBCs were incubated for 5 min with DAPI (0.1 μg/ml). iRBCs were imaged using a Leica confocal microscope. Acquired IFA images were processed by Image J software.

### Ghost preparation of taxol arrested P. falciparum infected RBCs

iRBC ghost was prepared as described in Das et al., 2012 (11). Briefly, taxol arrested iRBCs / iRBCs at about 6-7% parasitemia was pelleted at 500x *g* for 5 min and washed with cRPMI once. iRBC pellet was resuspended in 0.1% Saponin (Sigma S7900) and protease inhibitor cocktail (Sigma) and 1 mM PMSF in PBS, pH 7.4 for15 min at 37°C. The sample was then centrifuged for 10 min at 10,000x *g* at 4°C to get the parasite pellet and it was stored at -80°C. About 65–75% of the opaque supernatant (ghost and cytosol fraction) was gently separated to avoid cross contamination due to parasite pellet. This supernatant fraction was pelleted at 20,000x *g* for 2h at 4°C, washed twice with 1xPBS, pH 7.4, and stored at -80°C as IE ghost. After ghost precipitation, the supernatant (iRBC cytosol) was stored at -80°C for subsequent analysis.

### Immunoprecipitation assay (IP)

IP was performed as mentioned in Das et al., 2012 (11). Briefly, *P. falciparum* (Pf) parasite pellets were suspended in 200 µl non-denaturing lysis buffer (20 mM Tris HCl pH 8.0, 137 mM NaCl, 10% glycerol, 1% NP-40, 2 mM EDTA) in the presence of protease inhibitor cocktail (Sigma) in ice for 15 min. Cells were briefly sonicated (Branson Sonifier) for 1 min, and centrifuged at 15,000 *g* at 4°C. The supernatant was collected, protein was estimated using BCA (TaKaRa), and 100 µg protein was incubated with 20 µl packed protein G-sepharose beads (Sigma P3296) at 4°C for 1 hr for preclearing. Packed Protein G sepharose beads (20 µl) was washed repeatedly and the protein content of the pre-cleared lysate was estimated. For 3-4 µg protein lysate, 6-8 µg of anti-P2 antibody or control pre-immune serum was added. The protein-antibody solutions were incubated at 4°C for 6h on a rotary shaker, followed by incubation with 20 µl of packed Protein G-sepharose beads at 4°C for 2 hrs. Subsequently, the beads were centrifuged at 500 *g* and washed 6 times with lysis buffer. To the beads, SDS-PAGE loading buffer was added and boiled for 5 min followed by centrifugation at 15,000 *g* for 15 min at room temperature. The supernatant was loaded on SDS-PAGE for immunoblot.

### Immunoblotting

Recombinant PfP2 protein (2µg) under non-reduced and reduced conditions were run in SDS-PAGE. On the other hand, saponin freed parasite pellets were lysed in RIPA buffer (20 mM Tris HCl (pH 7.5), 150 mM NaCl, 1 mM Na_2_ EDTA, 1 mM EGTA, 1% NP-40, 1% Na-deoxycholate, 2.5 mM Na-pyrophosphate, 1 mM β-glycerophosphate) containing protease inhibitors (Sigma) by a brief sonication at 4°C or parasite pellet was directly lysed in SDS containing laemmli loading buffer in some experiments. The lysates were centrifuged at 15,000x *g* at 4°C for 30 min and the supernatant was used for the immunoblots. Before loading, the protein was mixed with gel loading buffer (50 mM Tris HCl pH 6.8, 10 mM DTT/100 mM β-Mercaptoethanol, 2% SDS, 0.1% bromophenol blue, 10% glycerol) and heated at 90°C for 10 min. Samples were resolved on 12% SDS–PAGE and proteins were transferred to methanol-activated polyvinylidene fluoride (PVDF) membrane (Millipore) using anode buffer (25 mM Tris HCl pH 10.4, glycine, 10% Methanol) and wet transferred for 2h at 4°C. PVDF membrane was blocked with 5% skimmed milk powder in 1x PBS overnight and probed with specific antibodies. Primary antibody dilution was made in 1x PBS containing Tween-20 (0.2%) and incubated with the membrane for 3 h at room temperature on a rocker. Primary antibody binding was detected by appropriate secondary antibodies conjugated to horseradish peroxidase (HRP) (AbCam). Dilution of secondary antibody was made in 1x PBS Tween-20 (0.2%). After each incubation, the membrane was washed with 1x PBS-Tween-20 (0.2%) for 5 min at least 5–6 times. The immunoblots were developed using the chemiluminescent substrate (BioRad).

### DNA electroporation and transgenic parasite selection

For conditional knockdown of P2 protein, homology region 1 (HR1) and homology region 2 (HR2) were PCR amplified from the coding region and from the 3’UTR respectively with two sets of primers. For HR1, P2FPHR1 and P2RPHR1 and for HR2, P2FPHR2 and P2RPHR2. Both the homology regions were cloned into the pL6-3HA-glms-ribozyme vector one by one with sequencing of DNA. Standard transfection method, parasite selection using 5nM WR99210 and negative selection with 5FC and limiting dilution to select integrated parasites were followed as described in Lyko et al., 2012 (62); Prommana et al., 2013 (63) and Ito et al., 2017 (64)). For Conditional down regulation of P2 protein under the *glmS* ribozyme system, 3mM glucosamine (GlcN) was added to synchronous trophozoite-stage cultures Prommana et al., 2013. GlcN exposure was continued for up to 12-14h before washout / harvest for phenotype studies. Control experiments with GlcN added in the culture media revealed no measurable toxicity in wild-type parasites at up to a 4 mM concentration.

P2FPHR1: 5’ATGGCTATGAAATACGTTGCTG3’; P2RPHR1: 5’CATTTCTACACATTTCATTTACATAATTT-TAAAG3’, P2FPHR2: 5’CATTTAGCACATTTTCCGTTTTGG3’ P2RPHR2: 5’GACAATTTGTAAAA-GAAAACTTCCC3’

### Cloning expression and purification of rec. PfP2

P2 gene from Plasmodium falciparum 3D7 (plasmodb gene ID:PF3D7_0309600) genome was amplified using forward primer 5′-CCCC***GAATTC***ATGGCTATGAAATACGTTGCTG-3′ and reverse primer 5′-GGGG***CTCGAG***TTAACCAAATAAGGAAAATCCTAAGTC-3′. PCR amplified P2 gene and pET28a expression vector were restrict digested using EcoRI and XhoI and P2 gene was cloned in pET28a. Transformed colonies in kanamycin LB agar plate were checked for the P2 gene using PCR. The *Plasmodium falciparum* 3D7 ribosomal protein P2 (PfP2) was expressed in the *E. coli* BL21 (DE3) cells transformed with pET-28 vector carrying the P2 encoding gene with 6X His tags at both N’ and C’ terminals. The overnight grown bacterial seed culture, obtained after inoculating 250µl of the glycerol stock along with 25µl of kanamycin (50mg/ml stock) in 25 ml fresh LB broth, served to be the seed culture for large scale growth at 37°C in 500 ml LB containing flasks. The cultures reaching OD ∼ 0.6 were induced with 1mM IPTG and grown till 3.5h at 37°C. Bacterial cell pellet was obtained from high-speed centrifugation of induced cultures. To carry out P2 purification, the cell pellets were resuspended in the prepared cell lysis buffer (0.1% Triton-X, 20 mM Tris, 100 mM NaCl) and ProteoGuard EDTA-Free protease inhibitor cocktail (Takara) for 30 minutes followed by sonication in ice cold water for 40 minutes (75 amplitude; 15 cycles, each cycle on for 10 sec and off for 59 sec). The supernatant obtained after the centrifugation of the lysate at 9000 rpm for 25 minutes at 4°C was subjected to overnight binding with Hi-bind Ni-NTA agarose beads (Invitrogen). The beads were washed 4-5 times with wash buffer (20mM Tris, 100mM NaCl, 50mM Imidazole, pH-7.4) and the His-tagged P2 was finally eluted with elution buffer (20mM Tris, 100mM NaCl and 200mM Imidazole, pH-7.4). A single band of monomeric protein (15 kDa) was observed in reducing SDS PAGE gel. The protein concentration was found to be 0.9 µg/µl using BCA Protein Assay Kit (TaKaRa).

### Blue Native PAGE (BN-PAGE)

4 µg of the rec. PfP2 was kept for 3h incubation at 37°C. The incubated sample was visualised in a BN-PAGE gel. The BN-PAGE was performed with the running buffer composed of 25mM Tris,, pH: 8.0, 192mM Glycine and 0.005% CBB R-250 at a constant current of 60mA. The incubated sample was mixed with native dye composed of running buffer, CBB R-250 and glycerol and it was resolved in a 12% Native-PAGE.

### Urea SDS-PAGE

4M urea (sigma U5378) was added in resolving and stacking gel. Additionally, 4M urea was also mixed with sample loading dye and in the running buffer. Gel was run like SDS-PAGE and stained with commassie R-250.

### Size exclusion chromatography of rec. PfP2

1.5 mg of pure PfP2 incubated at 37°C for 3h and was subsequently loaded onto a Hi Load Superdex 75 16/600 gel filtration column (GE Healthcare) pre-equilibrated with 20mM Tris-HCl (pH 7.4) and 100mM NaCl. Different fractions were accordingly collected at different time points and lyophilized.

### BN-PAGE and Atomic Force Microscopy (AFM) of rec. PfP2 and LAFS treated PfP2 tetramer

Each of the lyophilized fractions were visualized in BN-PAGE and the one showing single tetramer protein band was subjected to Atomic Force Microscopy. 1 µg rec. PfP2 / LAFS treated rec. PfP2 tetramer was firstly diluted 4000x in sterile filtered (0.22 µm membrane filter) milli-Q water to adjust the final concentrations of buffer salts in nanomolar range. 6 µl from the diluted sample was placed at the centre of the mica sheet (Muscovite V-1 Mica sheet ASP Cat no. 71856-04) and air-dried in a close cabinet (Petri dish). The prepared sample was observed under AFM using 9 µ scanner and cantilever oscillated in Acoustic AC mode. The 2D images were visualised in PicoView 2.0 software.

### Treatment of rec. PfP2 / LAFS treated rec. PfP2 with gradient concentration of surfactants (<CMC-CMC->CMC)

Gradient concentrations of four different surfactants, sodium dodecyl sulphate (SDS) (Sigma), sodium deoxycholate (Millipore 264101), CHAPS buffer (Sigma C3023) and Triton X-100 (Sigma T8787) were prepared. The concentrations were in the range starting below CMC (4 concentrations), CMC (1 concentration) and above CMC (1 concentration). rec. PfP2 / LAFS treated rec. P2 was treated with all surfactant concentrations at 37°C for 3h and resolved in Blue Native PAGE after mixing with 10 µl of native loading dye. Blue NATIVE gel running conditions was followed as mentioned previously.

### Treatment of rec. PfP2 with LAFS

4 mg of Albumax (AlbuMAX II, Invitrogen) was dissolved in 1xPBS (pH 7.4) and passed through a 5 kDa membrane cutoff (Millipore). The cutoff filter was then centrifuged at 1449x g for 45 mins at 4°C. 500µl of the Lower Albumax Fraction Solution (LAFS) was obtained at the bottom of cutoff and was treated with 0.5 µg of rec. PfP2 and incubated for 3h at 37°C. The LAFS treated rec. PfP2 sample was purified after binding with Hi-Bind Ni-NTA beads (Invitrogen) for 4h followed by elution of the sample with 20mM Tris-HCl (pH 7.4), 100mM NaCl and 200mM imidazole buffer. The elute was confirmed to be a pure tetramer in BN-PAGE. The confirmed LAFS treated rec. PfP2 tetramer was subjected for various biochemical and biophysical experimentations.

### Effects of surfactant gradient on LAFS treated rec. PfP2

The LAFS treated rec. PfP2 tetramer was added to six different SDS gradient concentrations prepared as mentioned before and visualized on BN-PAGE.

### LC-MS of not reduced, not Iodoacetamide (IAA) treated in-gel trypsin digested LAFS treated rec.PfP2 tetramer

In order to perform LC-MS to detect serum components, albumax (4 mg) was mixed with 1 ml 1x PBS and the solution was put through a 5 kD a membrane cutoff and centrifuged at 1449 x*g* for around 3h. LAFS obtained at the bottom of the cutoff tube was collected. 2µg of rec. PfP2 was mixed with 500µL of LAFS and incubated for 3h at 37°C with occasional vortexing to facilitate binding. 200 µL of Ni NTA beads (Invitrogen) was incubated with P2+LAFS for 2h at 37°C. Beads were washed and P2 was eluted with elution Buffer. LAFS treated P2 tetramers were separated in 12% SDS PAGE. Tetramer band was excised and subjected for destaining using detraining solution (80mg of ammonium Bicarbonate (NH_4_HCO_3_ and 20 ml of Acetonitrile and 20 ml of ultrapure sterile water). Destained gel pieces were dried completely and subjected for tyrosine digestion. 50 µL of Trypsin (1µg/0.1ml) was added to each of the bands and digested overnight at 4°C. Entire trypsin solution was removed and each of the gel pieces was chopped up. 10 µL of 1% TFA was added and incubated for 5 minutes to stop further digestion by trypsin. After removing the 1% TFA, the samples were further resuspended in 50µL of 0.5% TFA in 5% Acetonitrile. Peptides were further cleaned by C-18 resin columns (Agilent). (Cleanup and elution of peptides via C-18 column was performed as per the company protocols). Finally peptides were eluted in 0.1% TFA in 70% ACN. Precautions were taken to ensure that there was no keratin contamination while handling and processing the samples. ACN and TFA were mass spec grade.

### MALDI-TOF of rec.PfP2 / LAFS treated rec. PfP2 tetramer

To determine the mass of rec. PfP2 / LAFS treated rec. PfP2 tetramer, 500ng of protein was resuspended in 50µL of 0.5% TFA in 5% Acetonitrile (ACN). The C18 resin beads (Agilent) were first washed with 50% ACN solution and then equilibrated with 0.5%TFA in 5% ACN solution. Samples were then loaded onto the resin beads for binding. Resins beads were washed with 0.5% TFA in 5% ACN and the flow through was discarded and finally the sample was eluted using elution buffer comprising 0.1% TFA in 70% ACN. Eluted protein sample was then subjected for MALDI-TOF analysis.

### Synthetic fatty acids and rec.PfP2 interactions

Sodium salt of synthetic fatty acids (Sigma/Combi-blocks) were sparingly soluble in 1xPBS. Fatty acids were dissolved well after heat treatment at 70°C. 10mM of each fatty acid was incubated with 0.5 μg of rec.PfP2 at 37°C for 3h with constant mixing. As a control, 0.5 μg of rec.PfP2 was incubated in 1xPBS at 37°C for 3h. Protein samples were separated by non-reducing / reducing SDS-PAGE as mentioned or by 4M urea containing SDS-PAGE. Gels were stained by commassie R-250. PA: Palmitic Acid (P0500-25G / QF-5127), LA: Linoleic Acid (QD-5089), SA: Stearic Acid (QE-6119), MA: Myristic Acid (QE-1596), OA: Oleic Acid (O1008 / QA-7825), PTA: Palmiteloic Acid (QH-5487), Mix: Mixture of all fatty acids.

### Statistical Analysis

Statistical calculation was done and plotted as mean ± S.E.M. Significance was calculated by unpaired Student’s *t*-test or one-way ANOVA. Significance was considered at P<0.05 or indicated values. Not significant was denoted as P=NS.

## Figure Legends

**Supplementary data fig. 1.**
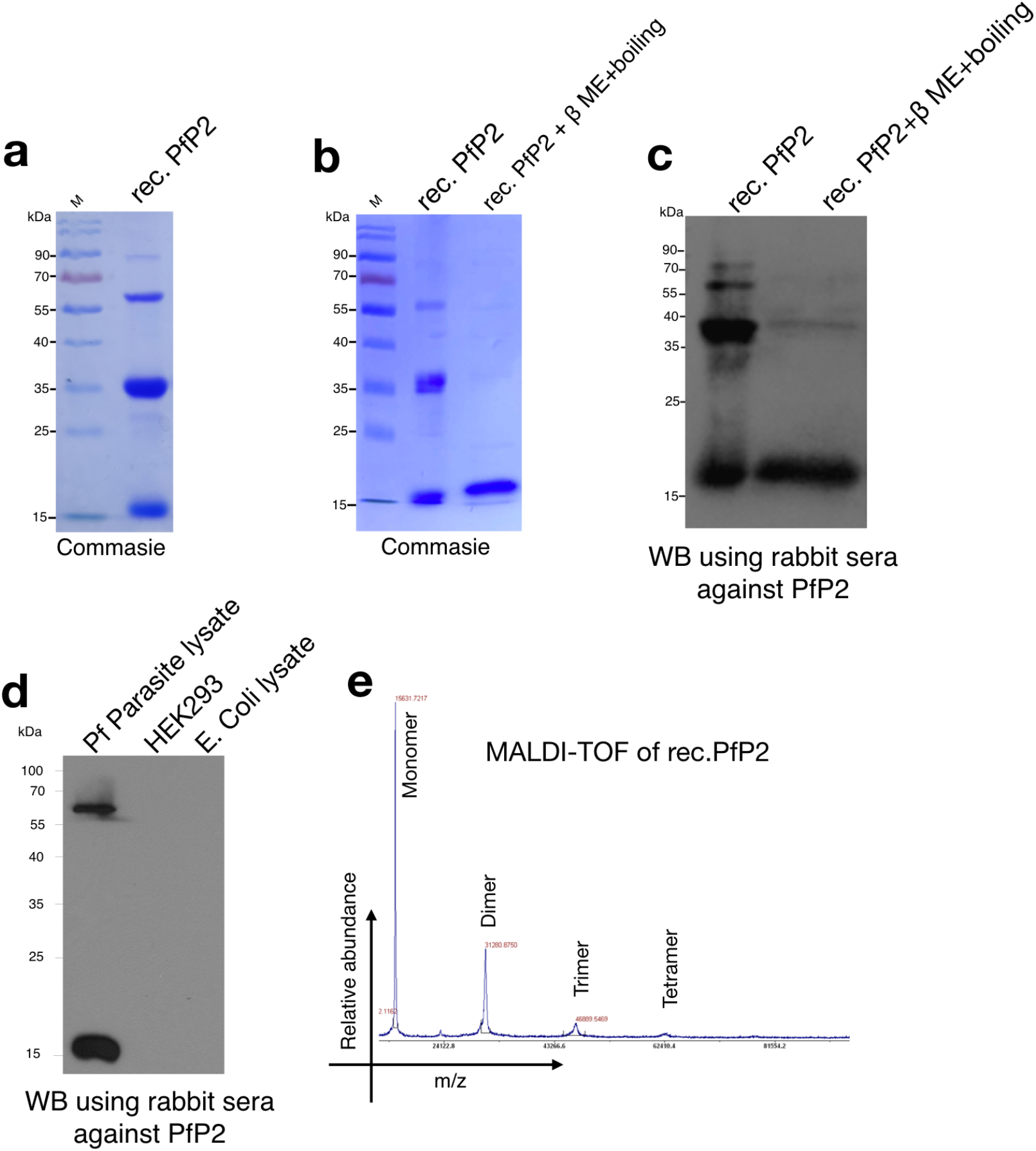
Under non-reducing condition rec.PfP2 exhibits monomeric, dimeric and tetrameric species. **a**. Non-reducing SDS-PAGE of affinity purified 6x His-tag rec. PfP2 shows different species of PfP2 protein. **b**. 1 μg of rec. PfP2 protein was separated in SDS-PAGE with/without reducing agent and boiling. **c**. Panel b was immunoblotted with rabbit anti-PfP2 polyclonal antibody. **d**. Immunoblot showing the specificity of rabbit anti-PfP2 polyclonal antibody 10k dilution. P. Falciparum 3D7 parasite lysate, HEK293 and E.coli lysates were used as control. 3 μg of total protein from lysate was separated in SDS-PAGE followed by immunoblotting using anti-PfP2 polyclonal antibody at 10k dilution. **e**. MALDI-TOF of purified 6x His-tag rec. PfP2 showing mass of different oligomeric species of PfP2 protein.

**Supplementary data fig. 2.**
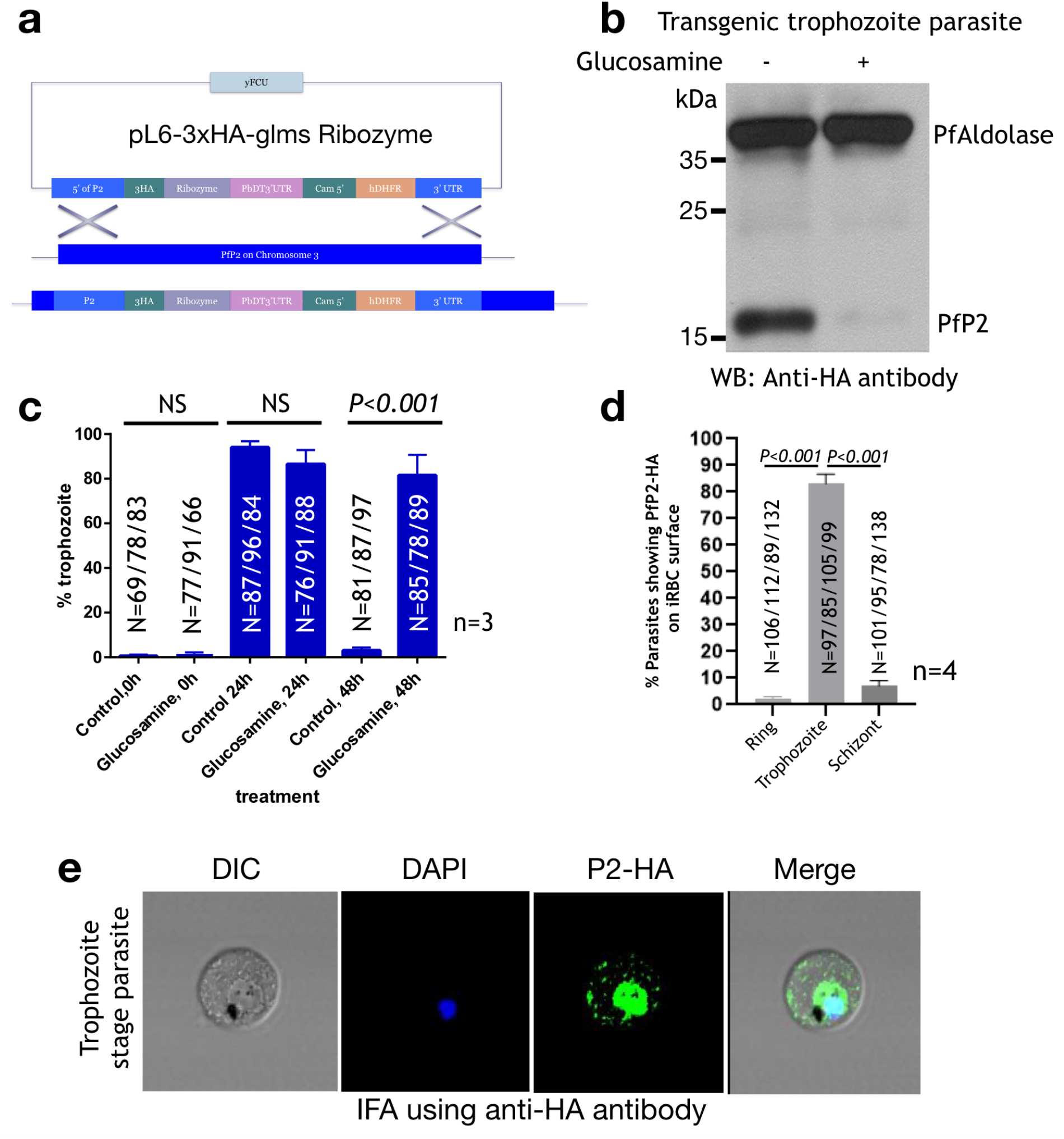
Down regulation of PfP2 resulted in arrest of parasite karyokinesis. **a**. Schematic of *PfP2* allelic exchange using pL6-glmS ribozyme system for conditional knockdown of P2 gene expression. **b**. Immunoblot of P. falciparum 3D7 transgenic trophozoite stage parasite cultured in the absence/presence of 3 mM glucosamine. 2 μg parasite lysate was separated in reducing SDS-PAGE and immunoblotted using anti-HA antibody. Parasite aldolase was used as loading control. **c**. Quantification of % trophozoite through Giemsa staining of parasites at different time points of parasite life cycle cultured in the absence/presence of 3 mM glucosamine. Statistical significance between two time points was not significant (NS) or significant as depicted P<0.001. **d**. Quantification of parasites showing PfP2-HA on infected RBC surface at three different stages of parasite development. Total number of parasites counted is depicted as ‘N’ at each parasite stage (n=4). **e**. Immunofluroscence assay (IFA) of transgenic P. falciparum 3D7 at trophozoite stage. P2 protein was detected using anti-HA antibody (green) and parasite nucleus was stained with DAPI (blue).

**Supplementary data fig. 3.**
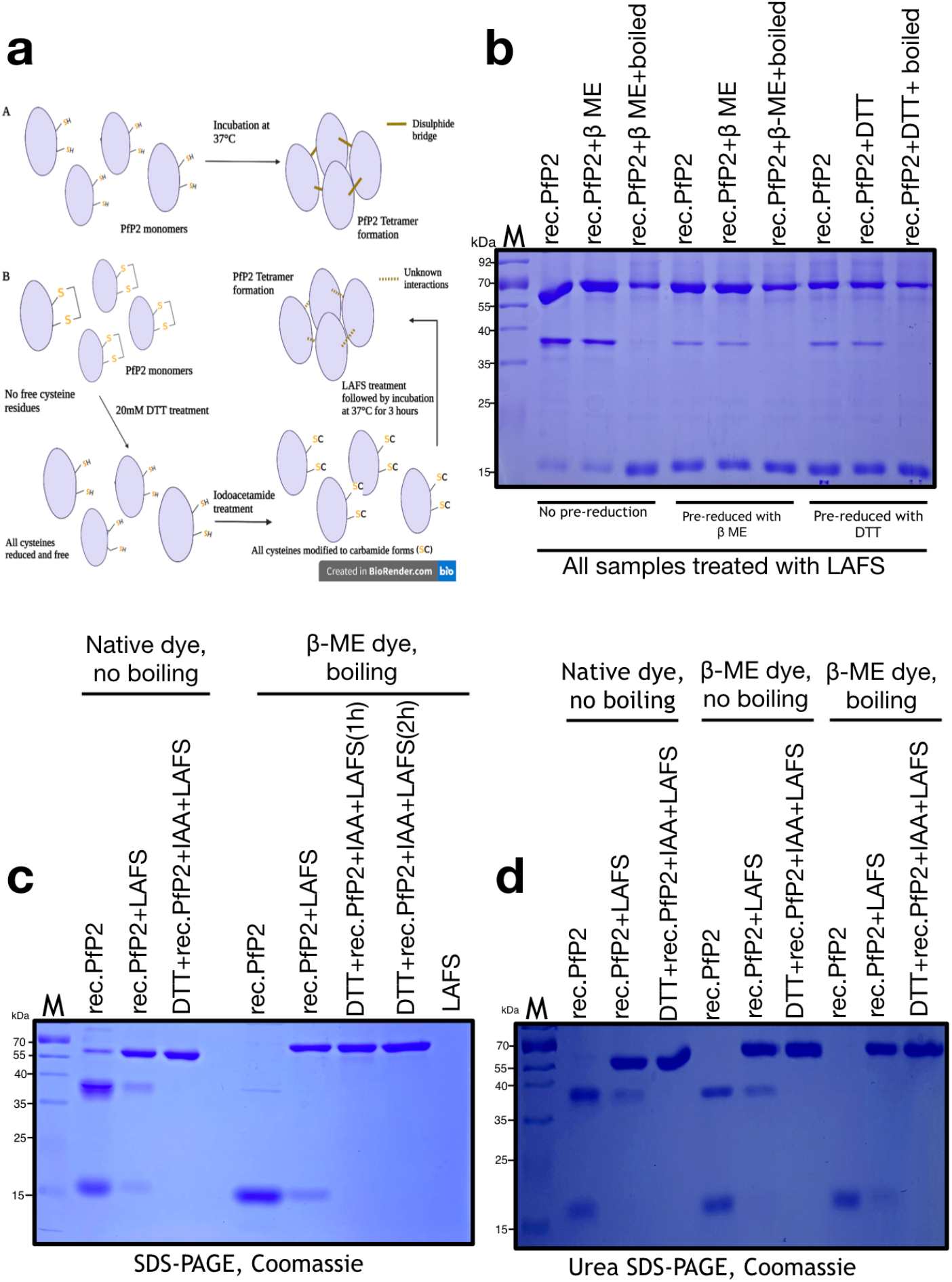
Denaturation resistance is not a consequence of NOS^40,41^ covalent linkage in LAFS treated rec.PfP2 tetramer. **a**. Cartoon explaining a hypothesis of the formation of denaturation resistant rec. PfP2 tetramer after LAFS treatment. **b**. 0.5 μg of rec. PfP2 protein was pre-reduced either with 100 mM β-ME or 20 mM DTT followed by LAFS treatment for 3h at 37°C. As a control, rec.PfP2 was not pre-reduce but treated with LAFS for 3h at 37°C. After treatment, all samples were mixed with either reducing or nonreducing dye and separated in SDS-PAGE. **c**. 0.5 μg of rec. PfP2 protein was pre-reduced with 20 mM DTT followed by Iodoacetamide treatment and then treated with LAFS for 1h and 2h. Treated rec.PfP2 protein was then separated in SDS-PAGE with native dye mixed with the samples and β-ME containing dye with boiling. As a control, only LAFS treated rec.PfP2 which was mixed with native dye and β-ME containing dye was used. Only LAFS was separated to check any protein contaminant from albumax. **d**. 0.5 μg of rec. PfP2 protein was pre-reduced with 20 mM DTT followed by Iodoacetamide treatment and then treated with LAFS for 3h. Treated rec.PfP2 protein was then separated in Urea (4M) continuing SDS-PAGE mixed with native dye, β-ME containing dye without boiling and β-ME containing dye with boiling. All the polyacrylamide gels were stained with coomassie R-250. All the gel images are the best representative and each type of experiment was repeated >3 times.

**Supplementary data fig. 4.**
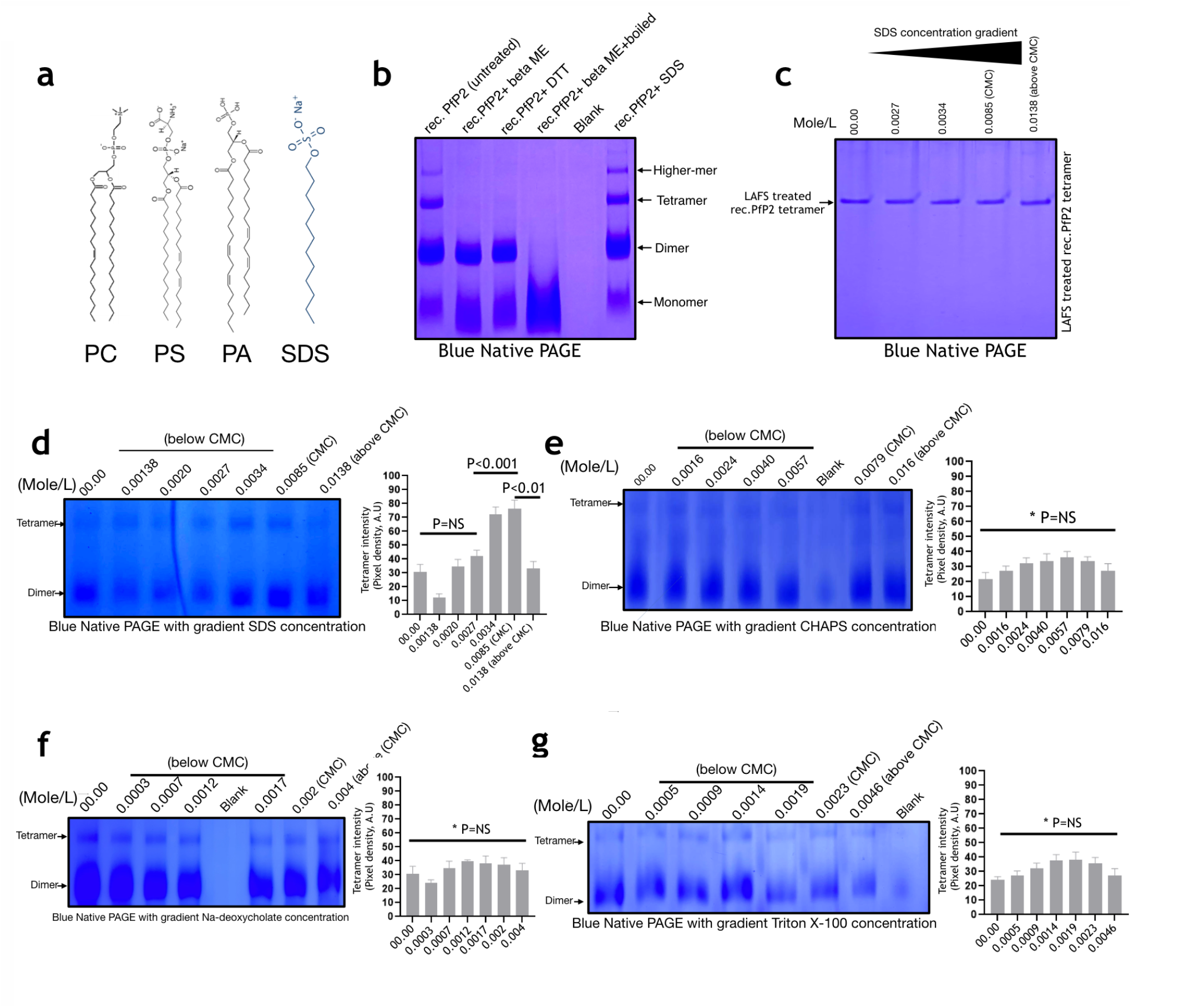
Acyl chain containing surfactants induce PfP2 tetramerization. **a**. Structures showing acyl chain containing phospholipids and sodium dodecyl sulphate (SDS). PC: Phosphatidylcholine, PS:Phosphatidylserine, PA: Phosphatidic Acid **b**. 3 μg of rec. PfP2 was separated in Blue Native PAGE under non-reduced (untreated), reduced (β-ME, DTT), reduced with boiling and after SDS treatment. At different lanes with different treatment, rec.PfP2 showing variation (reduced/enhanced) in oligomeric species. **c**. LAFS treated rec. PfP2 tetramer was treated with a gradient concentration of SDS (<CMC to >CMC). Treated rec. PfP2 tetramer was separated in Blue Native PAGE stained with commassie R250. **d**. 0.5 μg rec.PfP2 was treated with varying concentrations of SDS, <CMC to CMC and above CMC. Formation and degradation of P2 oligomeric spices were separated in Blue Native PAGE. The intensity of PfP2 tetrameric species with increasing concentration of SDS was quantified by Pixel density quantification using Image J software. The pixel density of tetramer from three independent blue native gel was averaged and plotted with an error bar (+/- SD). **e**. 0.5 μg rec.PfP2 was treated with varying concentration of CHAPS, <CMC to CMC and above CMC. Formation and degradation of P2 oligomeric spices were separated in Blue Native PAGE. Intensity of PfP2 tetrameric species with increasing concentrations of CHAPS was quantified by Pixel density quantification using Image J software. The pixel density of tetramer from three independent blue native gel was averaged and plotted with an error bar (+/- SD). **f**. 0.5 μg rec.PfP2 was treated with varying concentration of Na-deoxycholate, <CMC to CMC and above CMC. Formation and degradation of P2 oligomeric spices were separated in Blue Native PAGE. Intensity of PfP2 tetrameric species with increasing concentrations of Na-deoxycholate was quantified by Pixel density quantification using Image J software. The pixel density of tetramer from three independent blue native gel was averaged and plotted with an error bar (+/- SD). **g**. 0.5 μg rec.PfP2 was treated with varying concentration of TritonX-100, <CMC to CMC and above CMC. Formation and degradation of P2 oligomeric spices were separated in Blue Native PAGE. Intensity of PfP2 tetrameric species with increasing concentration of TritonX-100 was quantified by Pixel density quantification using Image J software. Pixel density of tetramer from three independent blue native gel was averaged and plotted with an error bar (+/- SD). Significant differences in the intensity of tetrameric spices in all the gels were depicted either non significant (P=NS) or significant (P<0.001; P<0.01).

**Supplementary data fig. 5.**
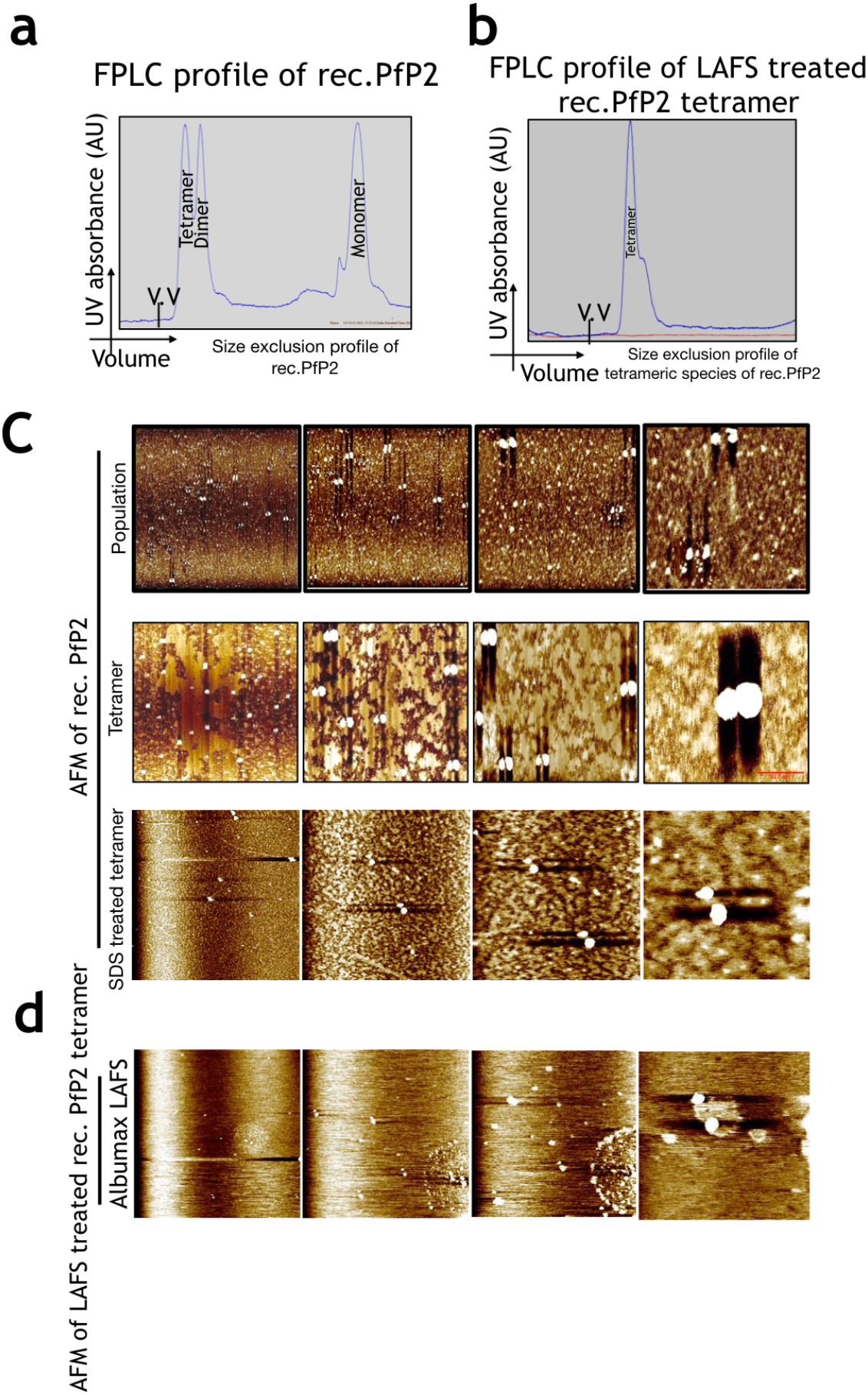
Unlike two monomers, two rec. PfP2 dimers assemble by S-S linkage to form rec. PfP2 tetramer. **a**. FPLC profile of 6x His-tag rec. PfP2 showing monomeric, dimeric and tetrameric species in superdex-75 gel filtration column. **b**. Gel filtration profile of LAFS treated rec.PfP2 in superdex-75 gel filtration column. Peak depicting tetrameric species of LAFS treated rec. PfP2. **c**. Atomic Force Microscopic (AFM) images of different species rec. PfP2, monomer, dimer and tetramer (population) and after SDS treatment (at CMC) at four different magnifications showing the structure of each species. **d**. AFM structure of albumax LAFS treated rec.PfP2 tetramer at four different magnifications. All AFM images are the best representative image of 5 independent experiments. The scale bar depicts the width of the PfP2 tetramer.

**Supplementary data fig. 6.**
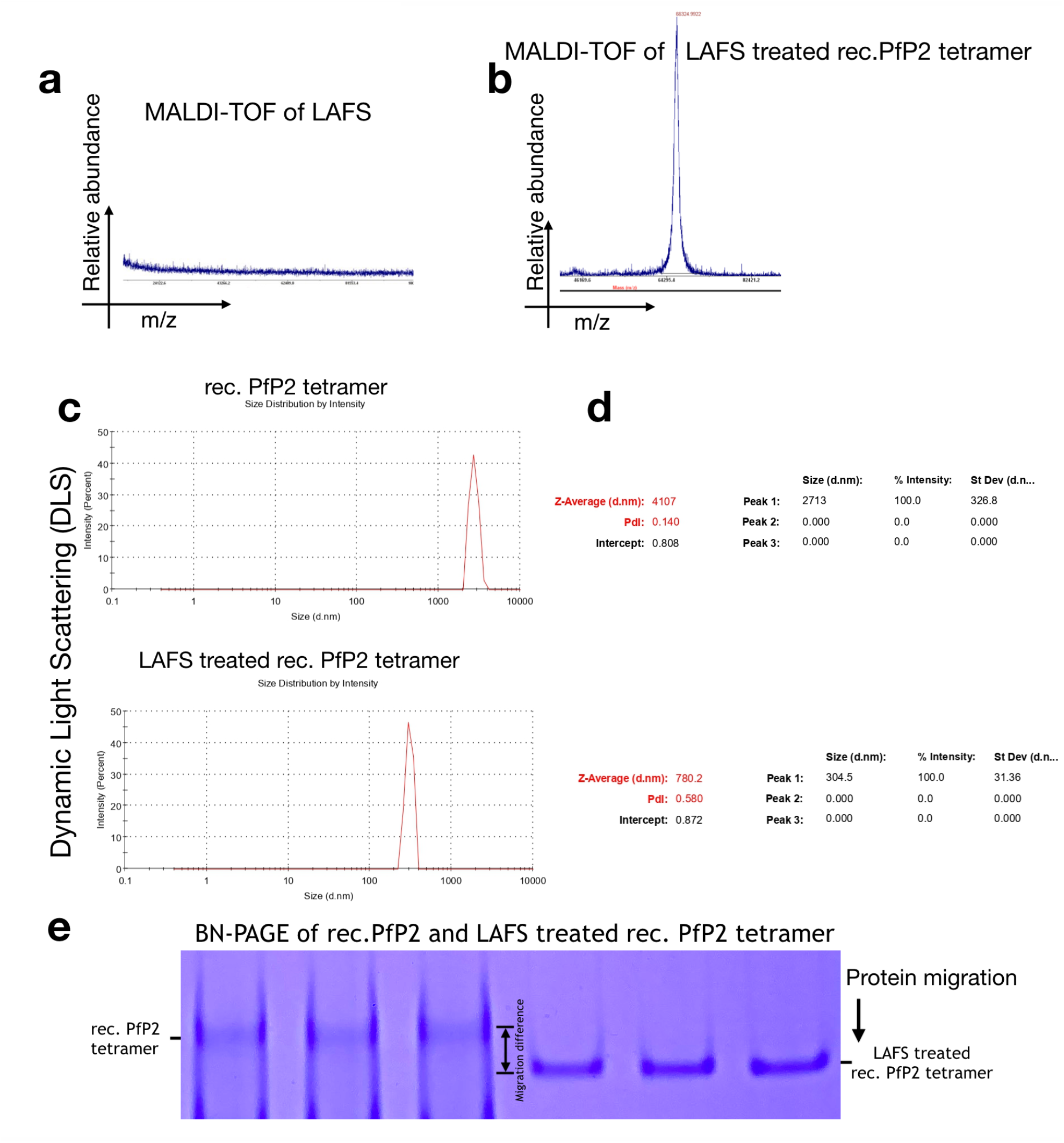
LAFS treated rec.PfP2 tetramer is smaller and compact in nature. **a**. MALDI-TOF analysis of LAFS. Y axis depicts relative abundance and X axis depicts m/z in the range from 20 kDa to 100 kDa. **b**. MALDI-TOF analysis of LAFS treated rec.PfP2 showing the mass of tetrameter as around ∼66 kDa. Y axis depicts relative abundance and X axis depicts m/z. **c. & d**. Dynamic Light Scattering (DLS) of purified rec.PfP2 tetramer and LAFS treated rec. PfP2 tetramer as depicted by graphical representation and d.nm. Differences in diameter between rec.PfP2 tetramer and LAFS treated rec.PfP2 tetramer is represented by both. **e**. Differences in migration between rec.PfP2 tetramer and LAFS treated rec.PfP2 tetramer is shown in Blue Native PAGE.

**Supplementary data fig. 7.**
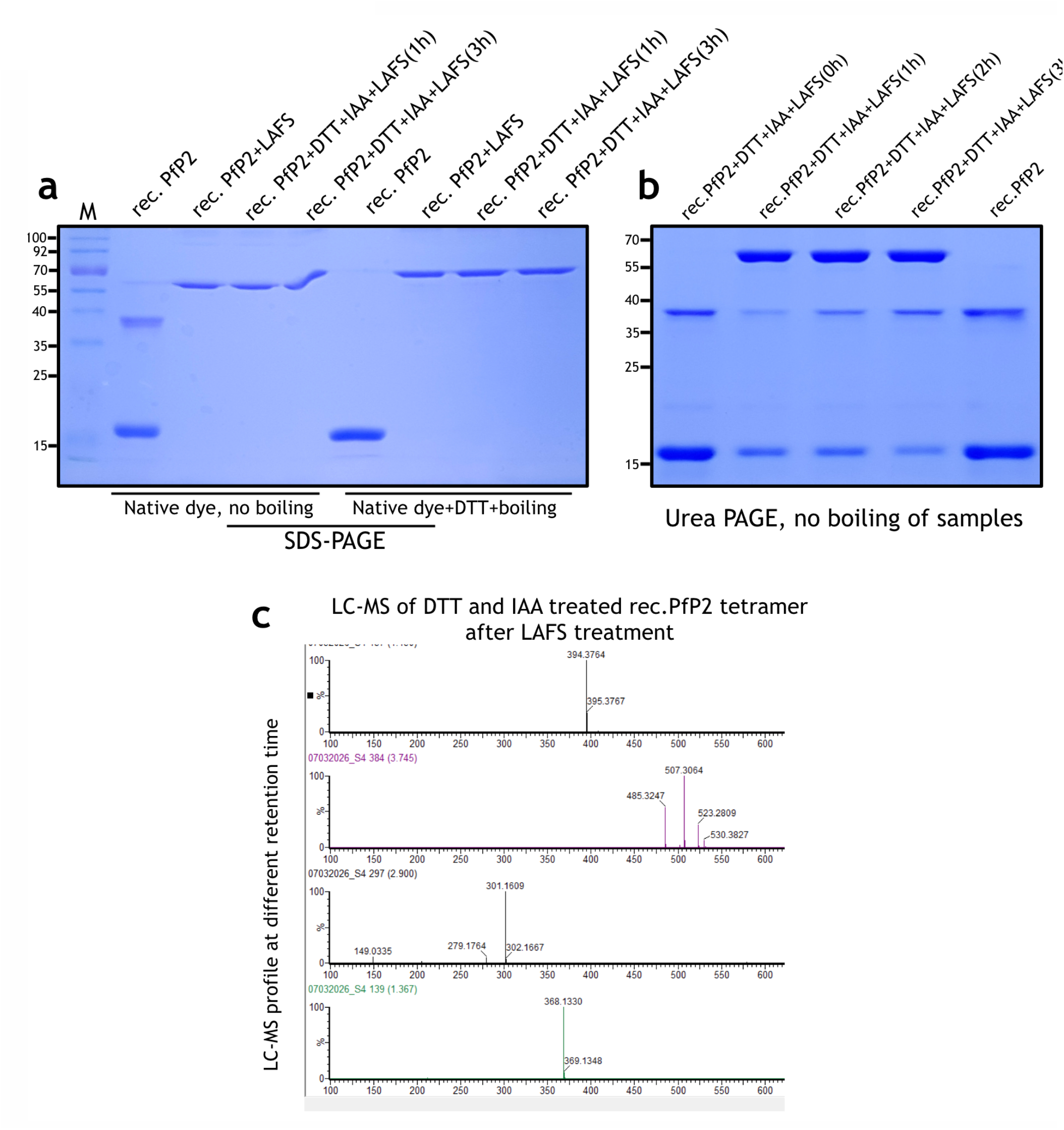
Cys-Cys redox switch is required to bind and release fatty acids. **a**. 0.5 μg of rec. PfP2 protein was pre-reduced with 20 mM DTT and then treated with Iodoacetamide (IAA). Treated rec.PfP2 was then treated with LAFS for varying times (1h and 3h). As a control, rec.PfP2 was only treated with 1xPBS and/or treated with LAFS for 3h at 37°C. Treated samples were separated in SDS-PAGE with native / reducing dye mixed with the samples with/without boiling. **b**. Samples from panel a were separated in 4M urea containing SDS-PAGE. **c**. LC-MS profile of rec.PfP2 tetramer which was initially DTT and IAA treated followed by LAFS treatment.

**Supplementary data fig. 8.**
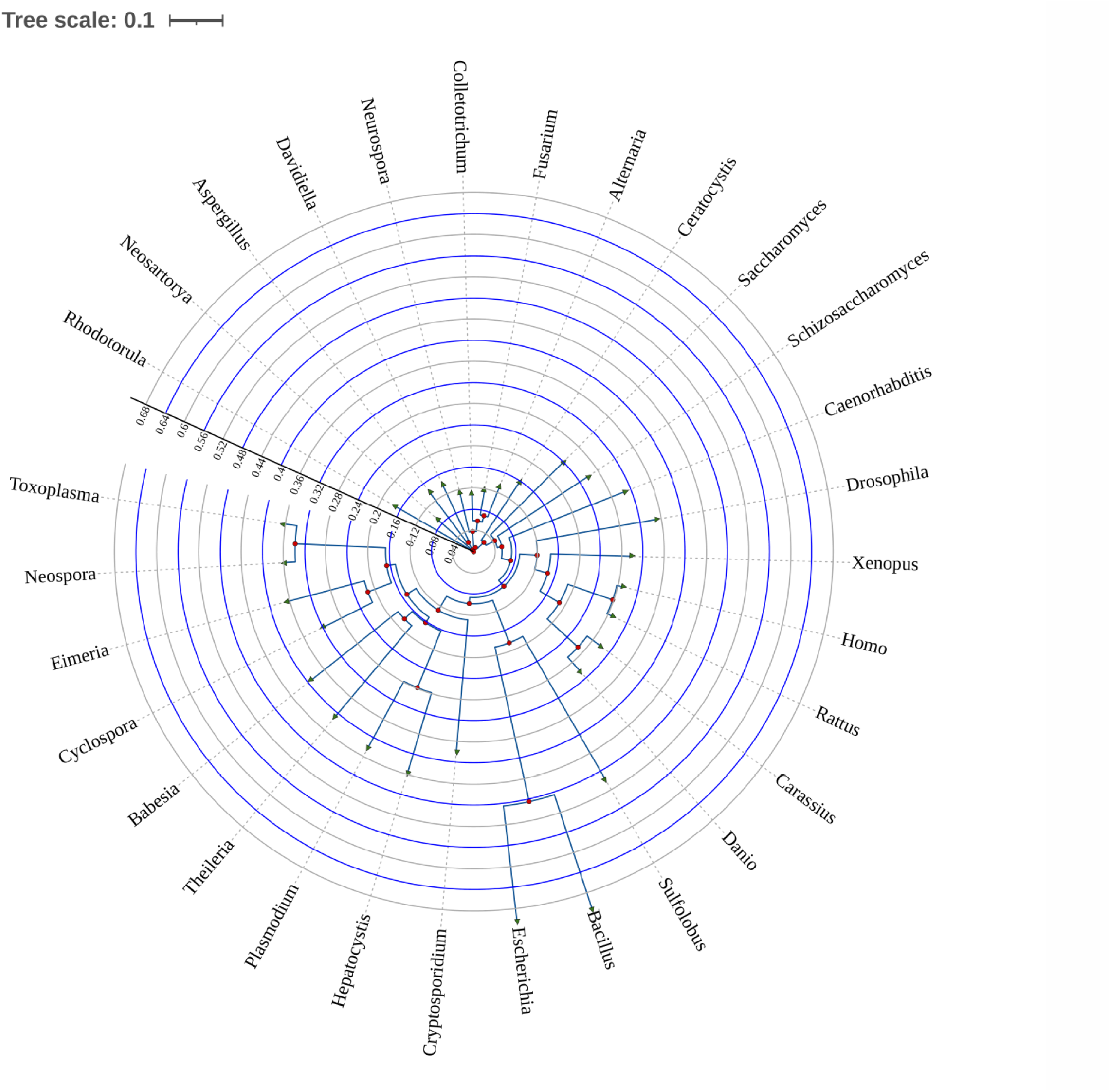
Phylogenetic analysis of P2 protein across living species. Phylogenetic analysis of ribosomal protein P2 sequence conservation and visualization in Tree of Life (circular mode). The ribosomal protein sequences have been referred to using the UniPROT database and blastp suite followed by alignment with the MAFFT tool (https://www.ebi.ac.uk/Tools/msa/mafft/). The internal tree scales (marked with two sets of lines, one in blue and the other in grey at intervals of 0.04 units) help to depict the lengths for each of the branches arising from the nodes. The divergence of organisms in terms of similarity in protein sequence could be delineated by comparing the branch lengths. The leaf nodes are marked with symbol ‘▷’(green in color) and internal nodes with ‘O’ (red in colour). The tree scale box is placed at the upper left corner.

## Additional Supplementary Informations

**Supplementary fig. 1.**
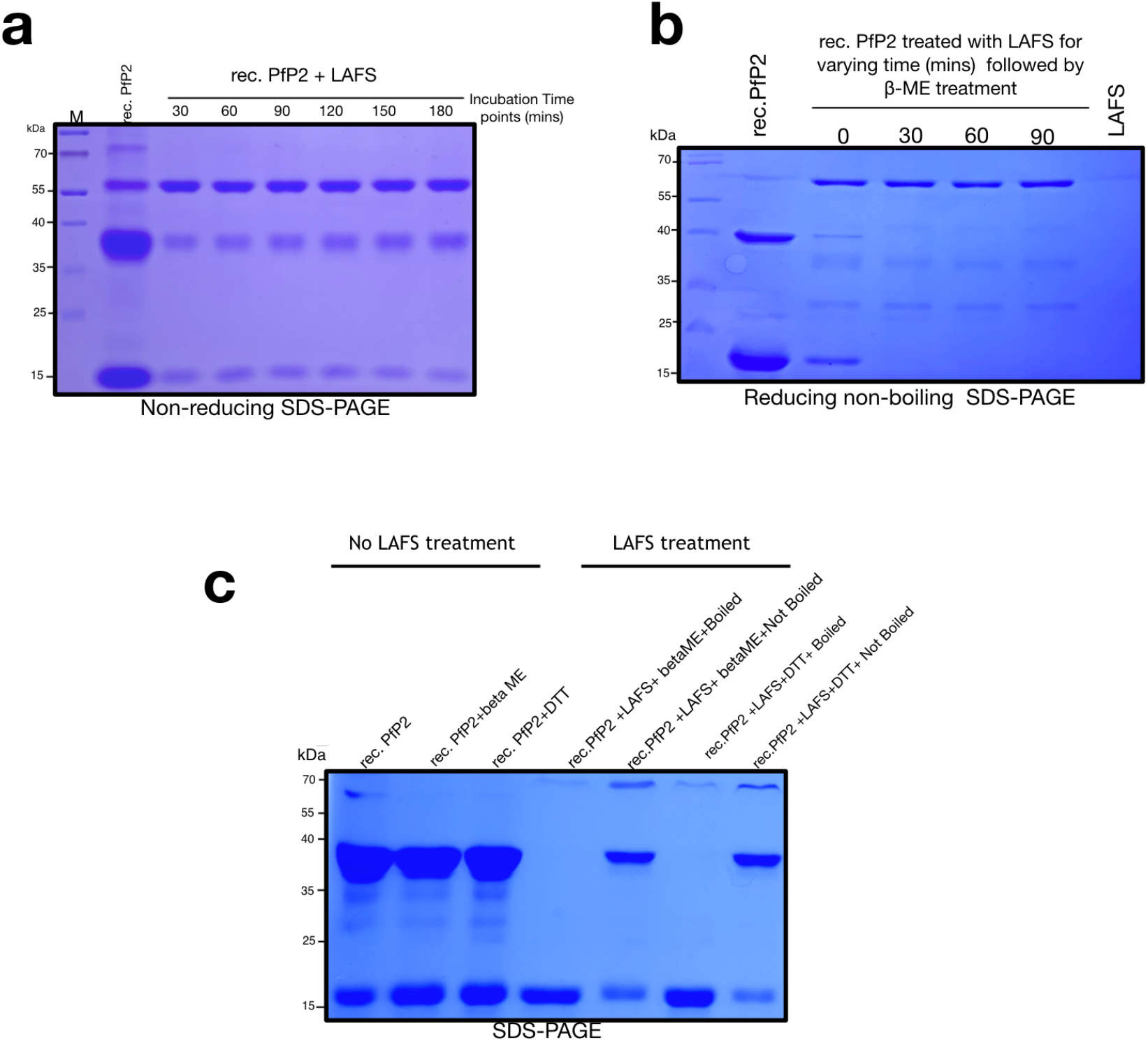
**a**. 0.7 μg of 6x His-tag rec. PfP2 was treated with LAFS at 37°C for varying time and separated in non-reducing SDS-PAGE. Rec. PfP2 in PBS for 3h at 37°C was used as a control. **b**. 0.5 μg of 6x His-tag rec. PfP2 was treated with LAFS at 37°C for varying time followed by 100 mM β-ME treatment and separation in reducing non-boiling SDS-PAGE. **c**. 0.7 μg of 6x His-tag rec. PfP2 was treated with LAFS at 37°C for 3h. Samples were reduced with 100 mM β-ME or 20 mM DTT with or without boiling and separated in SDS-PAGE. All gel images are the representation of > 3 independent experiments.

## Acknowledgements

We thank Prof. Sanjay A Desai from NIAID, NIH for providing pL6-HA-glms-ribozyme construct. We are thankful to MR4 BEI resources for Plasmodium parasite strains and other reagents. We thank Ashok Laboratory and Diagnostics, Kolkata, India for providing O^+^ human blood and human serum. We also thank Bioklone Biotech Pvt. Ltd, Chennai, India for custom made antibody synthesis.

## Authors contributions

SD: Conceptualization, designing experiments, performed experiments (parasite transfection, culture related work), data analysis, figure preparation, manuscript writing. KM: performed biochemical and biophysical experiments. AB: performed mass spectrometry. BR: cloning, biochemical and biophysical experiments.

